# Motile Living Biobots Self-Construct from Adult Human Somatic Progenitor Seed Cells

**DOI:** 10.1101/2022.08.04.502707

**Authors:** Gizem Gumuskaya, Pranjal Srivastava, Ben G. Cooper, Hannah Lesser, Ben Semegran, Simon Garnier, Michael Levin

**Author notes:** These authors contributed equally (Pranjal Srivastava, Ben G. Cooper). Corresponding Author: Dr. Michael Levin, Allen Discovery Center at Tufts University, Tufts University, 200 Boston Avenue, suite 4600, Medford, MA 02155-4243. Co-Corresponding Author: Gizem Gumuskaya, Allen Discovery Center at Tufts University, Tufts University, 200 Boston Avenue, suite 4600, Medford, MA 02155-4243.

## Abstract

Fundamental knowledge gaps exist with respect to the plasticity of cells from adult soma and the potential diversity of body shape and behavior in living constructs derived from such genetically wild-type cells. Here we introduce Anthrobots, a spheroid-shaped multicellular biological robot (biobot) platform with diameters ranging from 30 to 500 microns. Anthrobots have an inherent capacity for motility in aqueous environments, via cilia covering their surface. Each Anthrobot starts out as a single cell, derived from the adult human lung, and self-constructs into a multicellular motile biobot after having been cultured in extra cellular matrix for 2 weeks and transferred into a minimally viscous habitat. Anthrobots exhibit a wide range of behaviors with motility patterns ranging from tight loops to straight lines and speeds ranging from 5-50 microns/second. Our anatomical investigations reveal that this behavioral diversity is significantly correlated with their morphological diversity. Anthrobots can assume diverse morphologies from fully polarized to wholly ciliated bodies with spherical or ellipsoidal shapes, each correlating with a distinct movement type. Remarkably, as a function of these different movement types, Anthrobots were found to be capable of traversing live human tissues in various ways. Furthermore, Anthrobots were able to induce rapid repair of wounds in human neural cell sheets in vitro. By controlling microenvironmental cues in bulk, entirely novel structure, behavior, and biomedically-relevant capabilities can be discovered in morphogenetic processes without direct genetic editing or manual sculpting.

**Significance Statement:** We demonstrate that normal, non-genetically-modified human tracheal cells can be induced to form a new proto-organism - Anthrobots - which exhibit spontaneous behavior, swimming around in one of several patterns, demonstrating plasticity for novel form and function inherent in even elderly human somatic cells. Moreover, Anthrobots are able to traverse over cultured neurons, settling down and causing repair under them: the nerves knit together across the wound gap due to the presence of the Anthrobot. A patient’s own cells can be harnessed to make a motile biological robot that can traverse human tissue and induce repair. In the future, this platform can deliver pro-regenerative therapeutics for a range of biomedical applications that will not trigger rejection or require immune suppression.

## Introduction

What is the latent space of possible functional morphologies that cells, with a wild-type genome, can be coaxed to construct?^1^ This question drives at the heart of fundamental issues in evolutionary, developmental, cell, and synthetic biology, and has been taken up by a rapidly growing field focusing on building new kinds of active living structures: biobots.^2–6^ This emerging multidisciplinary effort to control the behavior of cellular collectives has garnered much excitement for two main reasons. First, because it offers the possibility of using engineering to reach outcomes that are too complex to micromanage directly, and hence promises to revolutionize efforts to produce complex tissues for clinical applications in regenerative medicine and beyond. Second, increased control over the morphology and behavior of cellular collectives could enable the development of self-constructing living structures by design with predictable and programmable functional properties and numerous practical uses, greatly extending the current abilities of traditional fabrication practices in diverse fields as robotics, architecture, sustainable construction, and even space exploration.

In the last decade, interest in developing biological structures *de novo* has seen a rapid surge ^7–24^. Among these efforts, a subset of functional biogenic assemblies gave rise to a special class of motile synthetic structures dubbed *biobots* ^25,26^. Early examples of biobots are hybrids between biological cells and inert chemical substances supporting them, such as gels or 3D-printed scaffolds.^27–31^ These efforts were followed by Xenobots, first fully-biological biobots created by sculpting or molding amphibian embryonic cells into multicellular structures that can spontaneously locomote without external pacing.^32–34^ Here we sought to address whether the capacity of genetically unaltered cells to generate a self-propelled, multicellular living structure in this way is unique to amphibian embryonic cells, and whether such a living structure can be built without needing to be individually sculpted or molded, but instead coaxed to *self-construct* from an initial seed cell, resulting in a high-throughput process wherein large numbers of biobots can be grown in parallel.

Here, we introduce novel, multicellular, fully-biological, self-constructing, motile living structures created out of human lung epithelium. We refer to them as Anthrobots, in light of their human origin and potential as a biorobotics platform,^2,5,6,35^ and we quantify their emergent, baseline properties as an essential background characterization of their native capacities from which future efforts to reprogram form and function will flow. Anthrobots self-construct in vitro, via a fully scalable method that requires no external form-giving machinery, manual sculpting, or embryonic tissues and produces swarms of biobots in parallel. They move via cilia-driven propulsion, living for 45-60 days. We quantitatively characterized the range of movement and morphological types, showing that their behaviors are strongly correlated with specific features of their anatomy. The ability of adult, somatic, human cells to form a novel functional anatomy, with unique behaviors, reveals that this plasticity is not restricted to amphibian or embryonic cell properties, and is a fundamental feature of wild-type cells that requires no direct genetic manipulation to unlock. Airway organoids with similar apical out tissue organization have very recently been shown using different protocols involving additional steps, ^36–38^ but these various alternative approaches have to date been used as organotypic cultures exclusively, and have yet to characterize the space of distinct movement or morphological types. Furthermore, Anthrobots exhibit a highly surprising behavior given their origin as static airway epithelium: they can traverse wounds in (human) neural tissue and induce their repair. Numerous in vitro and in vivo uses of such living machines can be envisioned, especially because they can now be made from the patient’s own cells.^39^

## Results

### Human bronchial epithelial cells self-construct into multicellular motile living architectures

To explore the self-organizing plasticity of morphogenesis without genomic change, we chose a cellular substrate in which such outcomes would be most surprising: adult, somatic, human airway tissues. To study and steer the in vitro morphogenesis of novel three dimensional tissues with motile appendages, we developed a novel protocol (Figure 1A) that builds upon the existing ability of human bronchial epithelial progenitor cells to form monoclonal spheroids (Figure 1-a.1) with cilia-lined lumina^40–43^ (i.e., apical-in configuration). We modified this process by manipulating the culture environment such that it now yields cilia-coated (i.e., apical-out) spheroids, which exhibit spontaneous locomotive ability.

**Figure 1.**
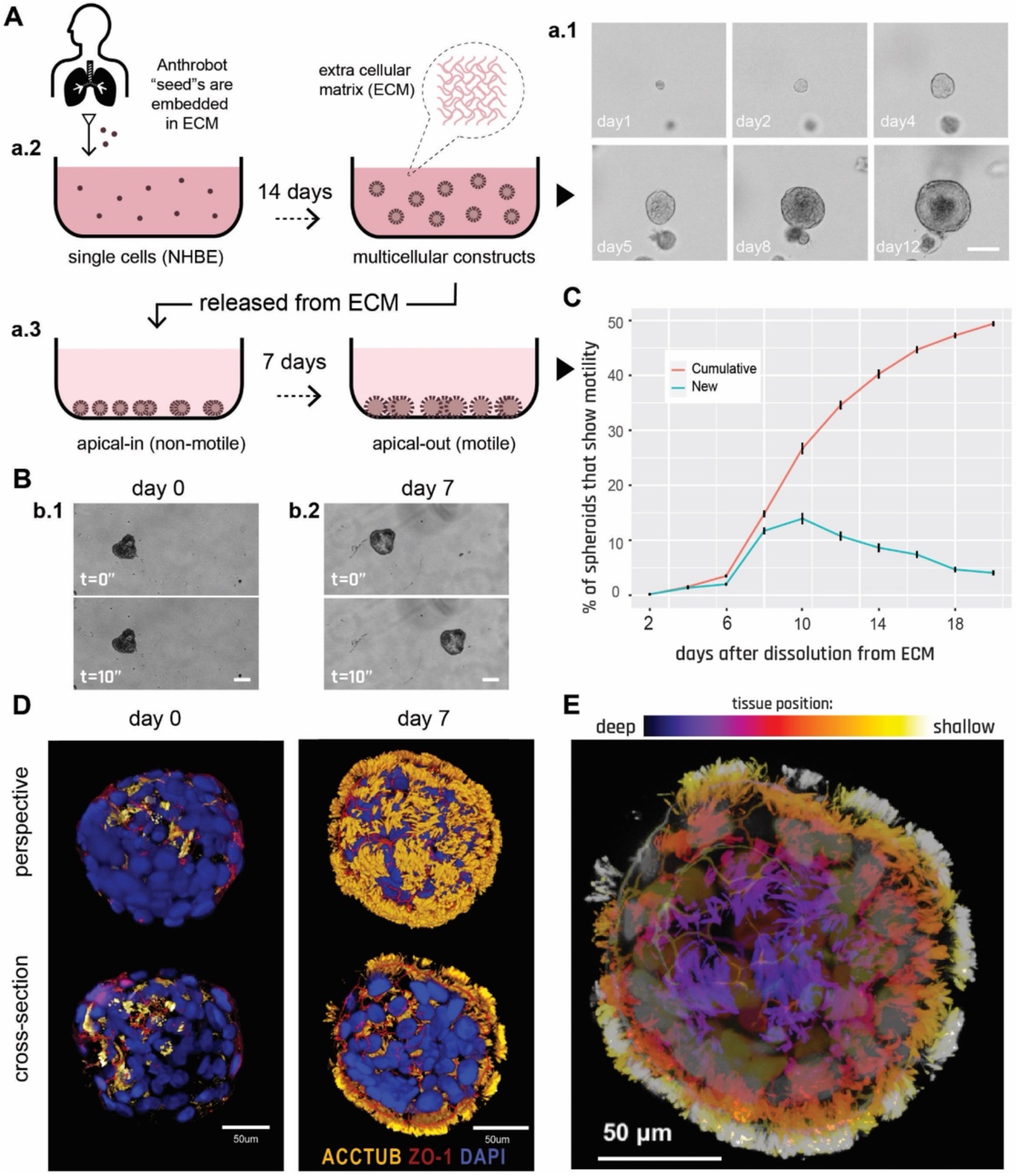
Human bronchial epithelial cells self-construct into multicellular motile living architectures. (A) Workflow for producing Anthrobots. NHBE cells’ apical-in to apical-out transition is facilitated by first culturing them in extra cellular matrix (ECM) under appropriate differentiation-inducing conditions), during which time apical-in spheroids self-construct from single cells (a.1), and upon the completion of this 14 day period (a-.2) by releasing mature spheroids from the ECM (a.3) and continuing to culture them in low-adhesive environment. Scalebar 50um. (B) Phase contrast images of an apical-in (b.1) and apical-out (b.2) spheroids, captured immediately after dissolution from ECM (day 0) and 7 days after dissolution (day 7), respectively. Day 0 spheroids show no motility, whereas day 7 spheroids show drastically increased motility. Scalebars 50um. (C) Percentage of cumulative and newly motile spheroids in the 3 weeks following dissolution. Out of the 2281 spheroids characterized total, around 50 % consistently showed no signs of motility (despite most having cilia) within this 3-week period and are referred as non-movers. (D) Immunostaining of day0 and day7 spheroids with a-tubulin (cilia marker), ZO-1 (tight junction marker), and DAPI (nuclear stain). Amount of multiciliated cells on the spheroid surface show a drastic increase by day 7. (E) A day 7 Anthrobot with depth information to show full cilia coverage. Immunostained with a-tubulin (cilia marker), ZO-1 (tight junction marker), and DAPI (nuclear stain). Colors represent tissue depth.

A key step in their construction, in order to obtain significant translocation, is the induction of cilia to face outward. Given that cilia naturally localize into the lumen due to the basal cells’ interaction with the surrounding high-viscosity matrix, we hypothesized that changing the culture environment to a lower viscosity level (e.g., water-based media instead of gel-based matrix) may trigger the basal layer cells to migrate inward and allow the apical layer to take their place on the spheroid cortex.^44^ Thus, to trigger apicobasal polarity switching, we first grew airway organoids embedded in Matrigel as reported previously^45^ (**Figure** 1-a.2), and then we dissolved the surrounding matrix and transferred the spheroids into a low-adhesive dish (Figure 1-a.3). In this new low viscosity environment, originally apical-in spheroids that show no motility on day 0 (Figure 1-b.1) became motile by day 7 (Figure 1-b.2).

To characterize the temporal dynamics of motility initiation, we periodically (every other day) counted the number of motile spheroids for 3 weeks following dissolution and observed a sigmoidal motility profile with peak change in motility on day 10 (Figure 1C). We next confirmed that this drastic change in motility occurred as a result of a morphological reorganization event exposing cilia on the cortex (Figure 1D). We immunostained the spheroids on day 0 (pre-motility) and day 7 (post-motility) with DAPI and for the apical markers a-tubulin (cilia marker) and ZO1 (tight junction marker), revealing a drastic increase in external multi-ciliated cells on day 7 compared to day 0. Figure 1E shows the tissue organization within an approximately 50-micron depth of a typical Anthrobot.

### Anthrobots self-organize into discrete movement types

Despite their wild-type human genome and somatic origin, these self-motile constructs exhibited wide range of behaviors and an anatomy that differs from the species-specific body morphology. As with any new behavioral subject ^46^, a key task is to determine whether its major properties and activities are discrete characters or continuous ones.^47,48^ Thus, we quantitatively analyzed their range of behavior modes in timelapse videos of approximately 200 randomly-selected motile spheroids (**Figure** 2A, see supplemental videos 1-4) for 5 hours in groups of 4 or 5 Anthrobots, and extracted their movement trajectory coordinates. We then split up these 5 hour-long trajectories into 30 second periods to classify behavior with a higher degree of granularity and in an aggregate manner. To identify patterns within a potentially unlimited set of possible movements, we characterized these periods by how straight and/or circular they are as all possible trajectories can be explained together by these two properties. To this end, we used two main trajectory characterization metrics: straightness and gyration indices (see Materials and Methods for detailed description of how these indices are calculated) and plotted all viable trajectorial periods along these two indices (Figure 2B). We then ran an unsupervised clustering algorithm on this two-dimensional plot and observed four statistically distinct clusters to emerge (Figure 2C). Further investigation of these clusters reveals that each represents a distinct movement type: circular, linear, curvilinear and “eclectic” (Figure 2D). Further analysis of each cluster in terms of its homogeneity (measured by “average dissimilarity” index), which is a measure of the intra-cluster variation, and its size (measured by “% of observations”) was performed (Figure 2E), as well as a quantitative comparison between different clusters along the two major movement indices (Figure 2F). See Supplemental Material “F2-Metrics Breakdown for Different Movement Types” section for more information.

**Figure 2.**
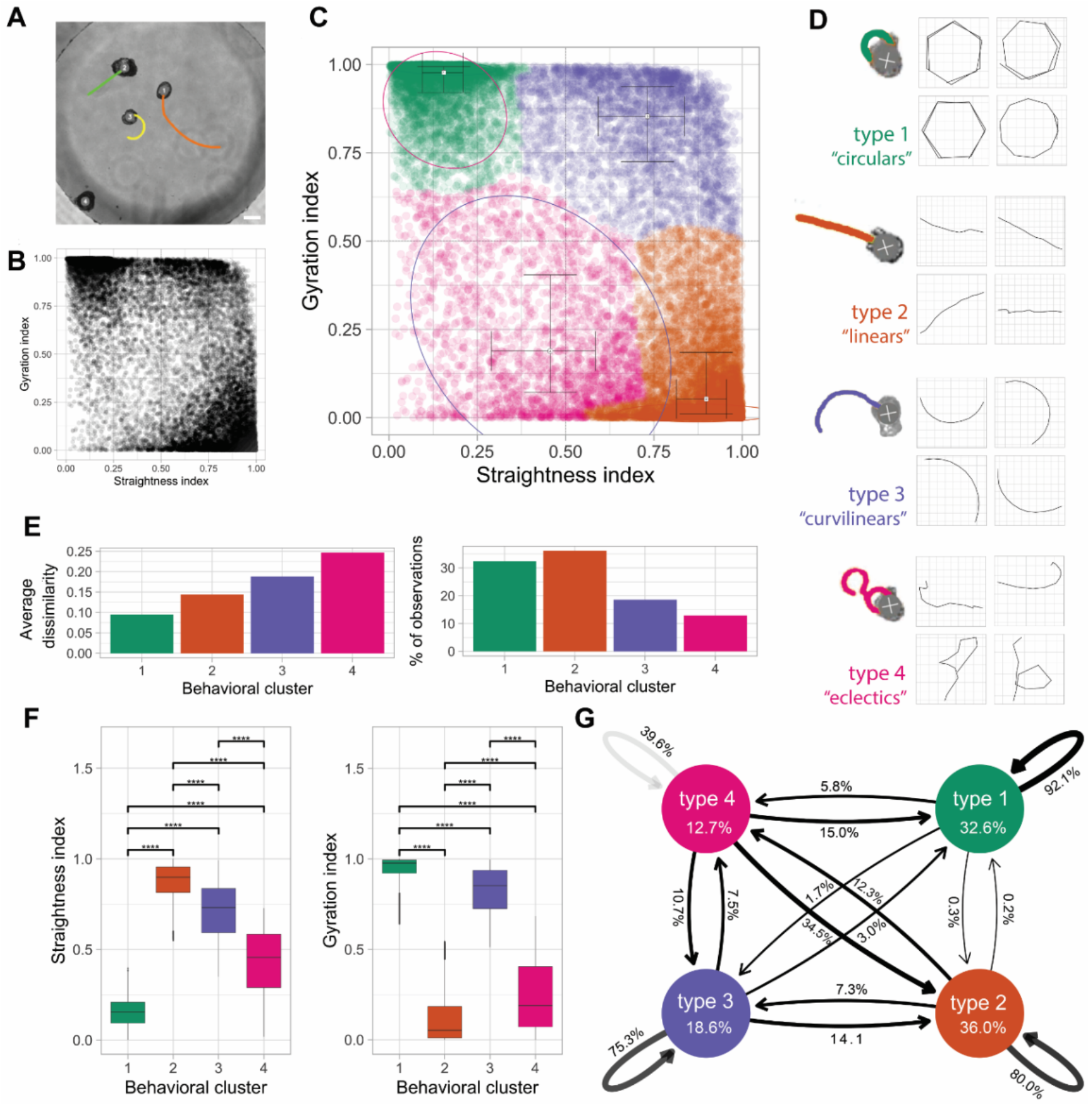
Anthrobots self-organize into discrete movement types. (A) Anthrobots display different movement types. Scalebar 100uM. (B) Distribution of all 30-second periods in the analysis plotted by their straightness and gyration indices, showing signs of clustering near three of the 4 corners of the plot. (C) Clustered scatter plot of all 30-second periods with centers of cluster marked and colored. (D) Prototypical examples from each cluster with 30-second sample trajectories. (E) Quantitative comparison of key characteristics of the four clusters in terms of intra-cluster homogeneity (“average dissimilarity”) and occurrence frequency (“% of observations”) which show that the largest clusters 1 and 2 have relatively low dissimilarity indicating these are the most consistent behavioral patterns. (F) Comparison of gyration and straightness indices for each cluster with significance levels indicated, showing that each cluster occupies a unique, quantifiable position in the sample space. (G) Markov chain showing state transitions between different clusters and the degree of commitment to a given behavior (persistence). See Supplemental Material “F2-PanelG” section for more information.

After having characterized each major movement type observed in Anthrobots, we next investigated the transition probabilities between each pair of behavior types. In order to estimate the stability of each trajectory and state transitions between different movement types, we used a Markov chain model shown on Figure 2G, which revealed the degree of commitment to a given behavior (persistence) and provided an ethogram of Anthrobot behavior. We observed that the most stable movement pattern for an Anthrobot is circular motion, followed by linear/curvilinear motion. The eclectics act more like an intermediate and over time, at least probabilistically, resolve into one of the 3 other categories. Therefore, we conclude that the vast majority of Anthrobot movements can be broken down into simpler, highly consistent patterns like linear, circular, curvilinear, with eclectics acting as a transient intermediary. The fact that Anthrobots exhibit movement types with high “consistency” and low rates of inter-type conversion (e.g., between circulars and linears) suggests that Anthrobots self-organize into discrete and stable movement types, each bot having a distinct motility fingerprint.

### Anthrobots self-organize into distinct morphological types

Having observed several distinct movement types, we next asked whether the range of Anthrobot morphologies was continuous or again composed of discrete categories.^49–52^ This question is important for both understanding the macro-scale rules of self-assembly, and for future efforts to control their functional properties. We hypothesized the primary parameters of this possible underlying morphological framework to be a function of the Anthrobots’ 3D shape and overall cilia distribution pattern, since Anthrobot motility is generated by cilia. Accordingly, we collected 3-dimensional structural data (**Figure** 3-a.1) from ~350 Anthrobots through ICC/IF and confocal microscopy, focusing on shape and cilia distribution pattern properties, and binarized these morphological features (Figure 3-a.2) to extract quantitative information on cilia and body boundaries. We then plotted this information for ~350 Anthrobots along eight different morphological characterization indices we developed, each quantifying a different aspect of the Anthrobot shape and cilia pattern. (Figure 3B, also see **Supplemental Figure** 2A). The shape-related indices among these eight formal morphological characterization indices included the ratio between the longest and shortest distance within a spheroid (i.e., “aspect”), longest distance within a spheroid (i.e., “max radius”), how invaginating or protruding the spheroid surface is (i.e., “shape smoothness”); the cilia-related indices included the total area covered by cilia signal on a spheroid surface (“cilia points”), cilia signal per unit area on a spheroid surface (“cilia points/area”), proximity of current cilia point distribution to a complete random uniform distribution (“cilia distribution homogeneity”), how clustered the cilia are on a spheroid surface (“polarity”), and how many free-floating cilia points there are that are not a part of a cluster (“noise points”). See Materials and Methods section for more details on how these morphological indices were calculated.

**Figure 3.**
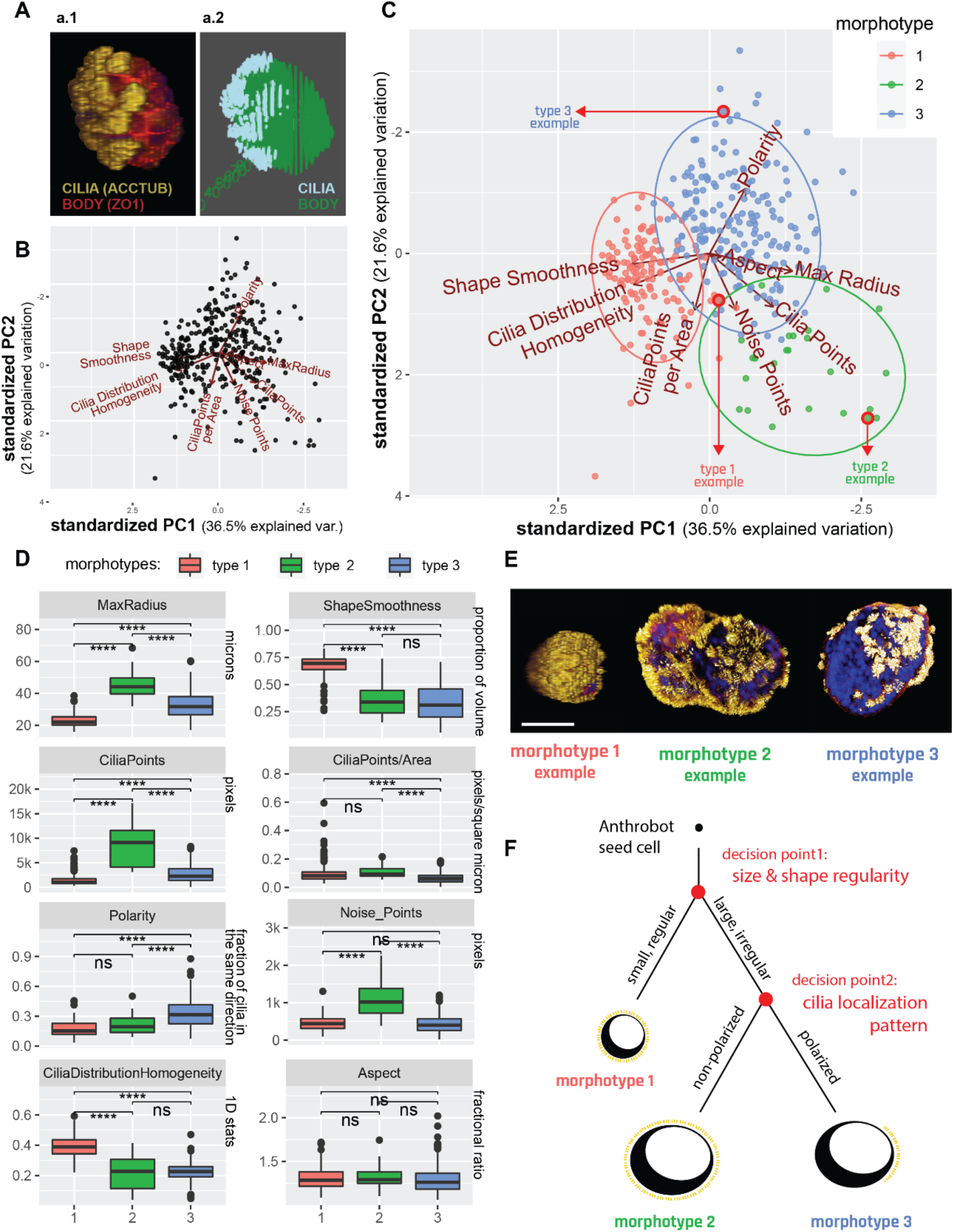
Anthrobots self-organize into distinct morphological types. (A) Anthrobot immunocytochemistry enables morphological classification pipeline. (a1) Sample immunological stain for cilia (acctub, i.e., alpha tubulin) and body mesh formed by the tight junctions (ZO1) (a2) and its binarized counterpart, showing the Anthrobot body boundaries and cilia localization on the body, represented by a mesh. (B) Binarized body and cilia information from 350 Anthrobots plotted along 8 morphological indices on a PCA cloud. (C) PCA clustered with the unsupervised Ward.D2 method with three morphotypical clusters emerging. Red arrows point to specific examples featured in panel E, selected from the cluster edges for distinct representation. (D) Distinct morphotypes translate with significance to differences in real-life morphological metrics, characterized by 8 variables from which the PCA was computed. (E) Sample morphotype examples for Type 1, 2 and 3 chosen for their ability to best represent the cluster. Type 1 Anthrobots are small, regularly shaped, tightly and uniformly covered by cilia. Type 2 and 3 bots are larger, more irregularly shaped and have less tightly-knit cilia patterns, with type 3 bots featuring significantly more polarized cilia coverage. Scalebar 50uM. (F) Decision tree of Anthrobot morphogenesis with two major checkpoints as revealed by the PCA hierarchy: first decision point is size/shape (has equal impact), second decision point is cilia localization pattern.

Next, we performed a PCA followed by an unsupervised clustering algorithm on this 8-dimensional data cloud and observed the emergence of three statistically distinct clusters (Figure 3C), each representing a distinct morphological type (morphotype). Figure 3D shows a quantitative characterization of each cluster along 8 different morphological indices. This analysis revealed the following two characteristics to be the most important distinguishing factors (to an equal degree, both ranking the top place in PC1 contributions) between different morphotypes: the size of the Anthrobot (measured by “max radius”), and the uniformity of its shape (measured by “shape smoothness”). These two most distinguishing characteristics formally describe type 1 bots to be significantly smaller in size and smoother (spherical) in its volume, while type 2 bots to be the largest and least uniformly shaped, and type 3 to be somewhere in between the two both. (See Materials and Methods section for PC1 and PC2 contribution rankings of different indices used to uncover this hierarchy.)

At the second level of importance in distinguishing between these three morphotypes is a set of four indices (all ranking equally top place in PC2 contributions), pertaining to cilia characterization. While the first two of these indices characterize cilia count, i.e., the density of cilia per Anthrobot (measured by “cilia points), and the density of cilia per unit area of Anthrobot (measured by “cilia points/area”); the remaining two indices characterize the pattern in which these cilia are distributed: how tightly grouped the cilia are (measured by “polarity,”), and the number of “free-floating” ciliary patches that are not within a group (measured by “noise points”). These four indices together describe type 2 bots as being significantly more ciliated than type 1 and type 3 bots, and type 3 bots as having a significantly more polarized cilia distribution pattern (with the least amount of extra-cluster noise) in comparison to type 1 and type 2 bots (Figure 3D).

The third most important (scoring a second level rank in both PC1 and PC2) characteristic in distinguishing between the different morphotypes is a function of both the size/shape of the Anthrobot and the localization pattern of its cilia: the homogeneity of cilia distribution on the surface of the Anthrobot (measured by “cilia distribution homogeneity”). This local index is related to, but not directly anti-correlated with, the polarity index, because while cilia distribution homogeneity characterizes local neighborhood patterns, polarity (along with its supporting index “noise points”) characterizes the global (entire Anthrobot-level) cilia distribution. (See Materials and Methods for more information). In this way, we obtain both a local and a global view of the cilia distribution patterns at once and identify type 1 bots as both globally and locally homogeneous, type 2 bots as globally homogeneous but locally heterogeneous, and type 3 bots as both globally and locally heterogeneous with high degree of global polarization.

In summary, our morphological characterization pipeline suggest that Anthrobots self-organize into 3 major morphotypes (Figure 3E) and this relationship can be represented by a developmental decision tree shown on Figure 3F wherein the first “decision point” determines the Anthrobot size and shape. Accordingly, bots that are small and regularly shaped (morphotype 1) form one branch, whereas bots that are larger and more irregularly shaped (morphotypes 2 and 3) form the alternating branch. On this alternating branch a second decision point forms further downstream and determines Anthrobot cilia pattern. Anthrobots with a non-polarized cilia pattern form one branch (morphotype 2), and Anthrobots with a polarized cilia pattern form the other (morphotype 3).

Finally, one characteristic that does not seem to be changing in any significant way between these three morphotypes is the ratio between the longest and shortest distance within a spheroid (measured by “aspect”). Although the 3 morphotypes differ significantly in terms of the volumetric regularities (measured by the shape smoothness index) as explained above, their aspect ratios are statistically very similar.

### Distinct movement types and morphotypes are highly correlated

Having observed the emergence of several discrete types of movement (Figure 2) and morphology (Figure 3), we next decided to investigate whether there is a mapping between Anthrobots’ different movement types and morphotypes. To do this, we incorporated an additional level of movement-type information into the PCA analysis used for identifying the morphotypes as introduced in Figure 3. During the initial sample collection process for this analysis, we had been able to definitively distinguish between non-motile Anthrobots (non-movers) and motile Anthrobots (movers) as described in Figure 1C. To further represent the movement types observed within the mover population, we randomly sampled from the set of motile subjects, targeting 30 Anthrobots that translocated (*i.e., displacing movers*) to assign a movement type. Selected displacing movers were randomly collected from the two most orthogonal movement types, circulars and linears, in approximately equal proportions.

Next, we identified these non-mover and displacing mover Anthrobots within the PCA cloud presented in Figure 3C and assigned them this additional layer of information, i.e., movement type (**Figure** 4A) without changing anything else in our sample pool or analysis workflow. In result, 62% of non-movers were identified within morphotype 1 cluster (with the remaining 38% falling into the morphotype 2 cluster). A 100% of displacing bots were identified in morphotype clusters 2 and 3, with ~85% of linear bots being in cluster 2, and 88% of circular bots being in cluster 3. We have further computed the statistical significance of these overlaps (using the Fisher test, see Materials and Methods) and conclude that the non-movers, linears, and circulars significantly correspond with the morphotypes 1, 2 and 3, respectively.

**Figure 4.**
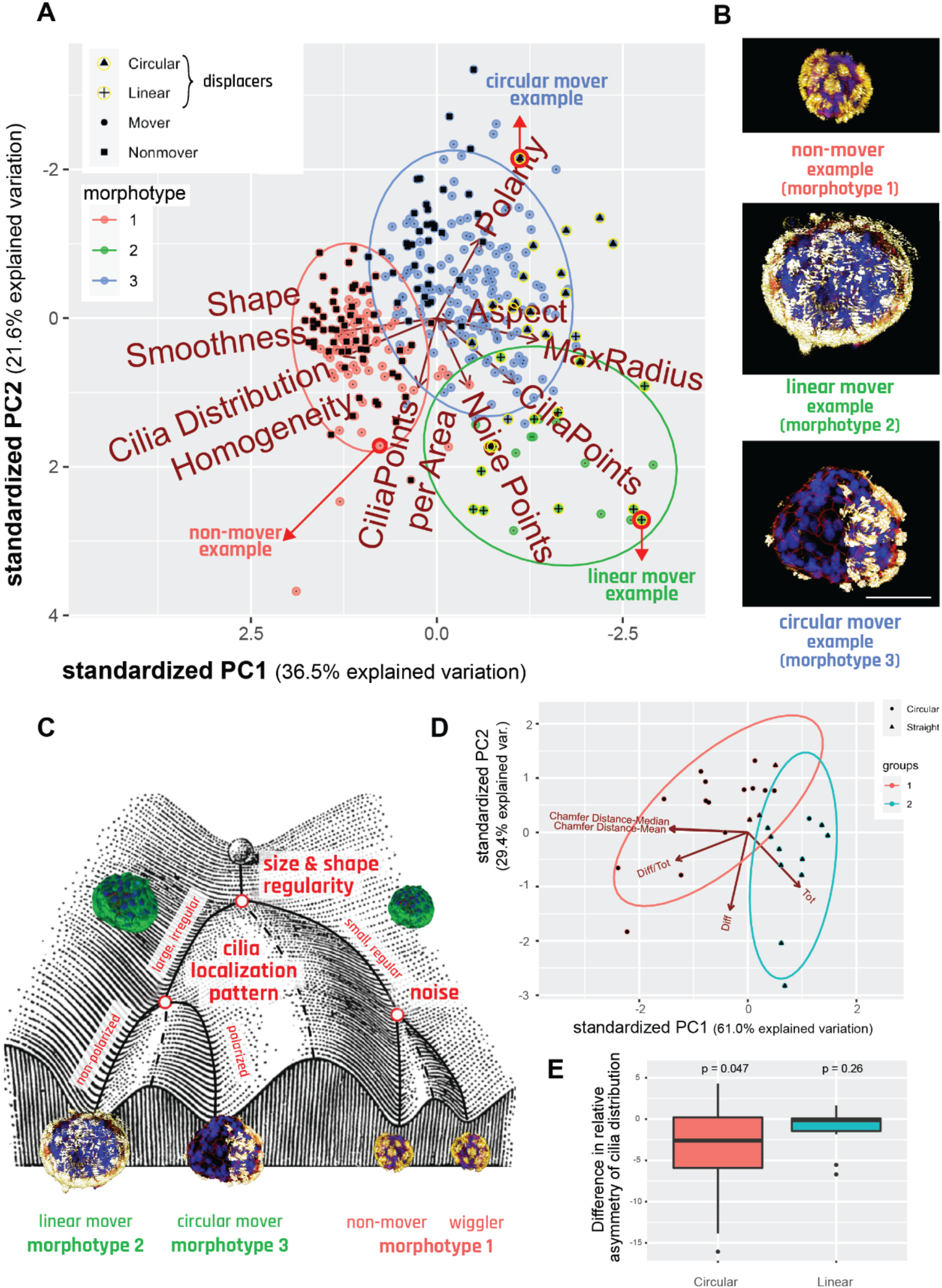
Distinct movement types and morphotypes are highly correlated. (A) PCA of 350 bots forming 3 morphotypical clusters, showing that there is significant overlap between these clusters and the separately marked Non-movers, linears and circulars. Red arrows point to specific examples featured in panel B, selected from the cluster edges for distinct representation. (B) Sample morphotype examples from each cluster, corresponding to Cluster 1, 2 and 3 and Nonmover, Linear, and Circular, respectively chosen for their ability to best represent the morphotype vs movement type mapping. Scalebar 50uM. (C) Waddington landscape illustrating the logic of determination of bot behavior and their relation to morphotypical indices with end behavioral products, as well as the potential states possible at each level of bifurcation of the bots’ development. (Waddington Landscape image modified from J. Ferrell, 2012.) (D) PCA and unsupervised clustering showing the polarization among linear and circular bots in respect to bilateral symmetry metrics. (E) Difference in asymmetry of cilia distribution between the movement axis and its 90-degree offset axis.

The fact that none of the displacing bots were identified within the morphotype 1 cluster suggests that the movers identified within this cluster (i.e., those morphotype cluster 1 data points which are not labeled as “non-movers”) are statistically likely to be *non-displacing movers*, displaying a stationary wiggling motion. Accordingly, we conclude that morphotype 1 bots are likely to assume either non-mover behavior or wiggler behavior. This may be attributable to their spherical shape with homogeneously distributed cilia where the propulsion forces generated by the ciliary motion are more prone to canceling each other out due to the radially symmetric spherical shape, resulting in little or no movement. Accordingly, inherent noise in the system (such as small imbalances in the cilia distribution on the spheroid surface or how the bot happened to be oriented in the plate) may be sufficient to have these bots generate small amounts of movement, causing them to wiggle, but not enough movement to become a displacing bot.

As a result of these analyses, we conclude that there is a statistically significant relationship between the Anthrobots’ developmental morphology and their behavior, and we show visual examples of this relationship with categorical examples on Figure 4B. We further represent this relationship by a decision tree in the form of a Waddington Landscape – a formalism often used to characterize cell- and body-level properties by mapping out the sequential logic of decision-points in transcriptional space or morphospace.^53–56^ Figure 4C shows the Waddington Landscape for the Anthrobot. The single cell at the top of the diagram represents the single cell that will develop into the multicellular Anthrobot. During this process of self-construction, the Anthrobot moves through the developmental landscape, negotiating certain points of morphological possibility to reach its final architecture. We conclude that the unique and spontaneous 3-D multicellular morphogenesis of adult airway cells into Anthrobots is consistent; the final form of the Anthrobot displays a degree of variability and exhibits discrete characters with easily recognizable primary features that also map on to phenotypic behavior.

### Anthrobots show bilateral asymmetry along movement axis

The above metrics all focused on the global structure of the bot. Next, we studied the local characteristics that connect the movement of bots to their morphology, by looking for a difference in bilateral symmetry, or lack thereof, between the two major types of displacing bots (linears and circulars) through symmetricity measurements across plane coincident with their direction of movement. One hypothesis is that Anthrobots have bilateral symmetry that underlies their ability to move in straight lines (as observed in many existing species ^57,58^ and even synthetic forms ^59^); this hypothesis predicts that Anthrobots with circular motion should have more asymmetry across their movement axis compared to other planes. This hypothesis was tested by running a PCA and unsupervised clustering algorithm on a point cloud quantifying Anthrobot cilia distribution patterns through four major bilateral symmetry-related measurements: total cilia points on a given bot (measured by “tot”), difference in number of cilia points between the two hemispheres (halves of the bot that are separated by the movement axis) of a given bot (measured by “diff”), this difference normalized by total cilia points (measured by “difftot”), and finally the bilateral symmetry index along the movement axis (measured by “Chamfer distance,” see methods for more details). Results of this analysis (Figure 4D) yielded two major clusters that in a statistically significant manner, each corresponding to one of the two major types of displacing bots (linears and circulars), with the group consisting predominantly of linears scoring significantly higher on the bilateral symmetry measurement (via Chamfer Distance axes, which inversely correlate with bilateral symmetry measurement). This result provides preliminary evidence in support of the hypothesis that Anthrobots with linear movement trajectories may have higher degrees of bilateral symmetry.

In order to further test this hypothesis, while also controlling for the globally homogeneous cilia distribution in linear bots posing a potential confounding factor for bilateral symmetry measurements, we compared the bilateral symmetricity of linear and circular bots against arbitrary axes other than the axis of movement. The initial hypothesis that linear Anthrobots may have higher degrees of bilateral symmetry compared to circular bots automatically suggests that for linear bots, we would expect there to be no other axis than the axis of movement along which the bilateral symmetry is higher; and for circular bots, we would expect there to be other axes than the axis of movement in respect to which the bilateral symmetry is higher. We tested this postulation by measuring linear and circular Anthrobots bilateral symmetry indices separately along each bot’s axis of movement vs its farthest rotated (i.e., 90-degree rotated) counterpart (as the control axis). As a result (Figure 4E), we indeed observed that while for the linear bots there exists no other axis in respect to which the bilateral symmetry is higher than that of the axis of movement, for the circular bots, there exists other axes than the axis of movement in respect to which the degree of bilateral symmetry is higher (p = 0.047). (See **Supplemental Figure 3** for comparison with other control axes that have rotational angle smaller than the farthest possible 90-degrees.) Taken together, these findings support our hypothesis that Anthrobots with distinct movement types have distinct local bilateral symmetry profiles, with linear bots showing higher bilateral symmetry. This suggests that these synthetic forms recapitulate a fundamental morphological property observed in many wild-type species.

### Anthrobots can traverse live tissues in vitro

One possible use of these living machines is to manipulate other tissues, in vitro or in vivo, in future biomedical or bioengineering applications. How will biobots react to environments different from those that their component cells face in their native configuration in vivo? Thus, Anthrobot behaviors need to be characterized outside of a bare culture dish context, and especially in environments that airway epithelia do not normally encounter. Having characterized their baseline movement and morphology, we wanted to assay this novel motile form for potentially useful behaviors and ways in which it may interact with other somatic tissues, especially sites of damage. Because we are interested in surprising examples of behaviors in such novel constructs, we sought to confront them with a scenario which would not be natural for these airway cells, either in vivo or in their evolutionary history. We decided to study the ability of Anthrobots to traverse live tissues that have been damaged, taking advantage of a common model system: the scratch assay in vitro.^60,61^ We produced 2-D confluent layers of human neurons derived from hiNSCs based on a previously established method,^62^ and introduced a wound of 200-400 microns by mechanically scratching away the neuron layer.

Anthrobots were placed within these “neuronal wound” environments in order to characterize their dynamics in this novel biological environment. Bots were allowed to freely move on their own and timelapse videos were recorded (**Figure** 5A, see **supplemental videos** 5, 6). These videos were then tracked, and specific indices were calculated from the tracked files (Figure 5B). Among these indices, we characterized the degree to which bots interact with the native tissue surrounding the wound (measured by “proportion of bot on tissue”), bots’ tendency to assume a circular motility profile (measured by “bot’s rotational tendency”), and bots’ displacement speed in traversing the wound (measured by “instantaneous velocity”). More specifically, we investigated the relationship between bots’ efficiency in traversing the wound as a function of the circularity of their movement pattern (Figure 5C). We observed a significant positive relationship (slope=1.5, p=0.01), confirming our baseline assumption that although circling bots are less efficient in forward motion, they are better at “exploring” the wound. We have further observed that the instantaneous velocity also had a significant positive relationship with bots’ efficiency in traversing the wound (slope=0.0082, p=0.031 (Figure 5D), presumably due to increased collisions with the tissue. Taken together, these data reveal that Anthrobots are capable of traversing damage sites in tissues, and that bots that have a higher rotational tendency and or higher speed end up exploring the wound better by covering a higher percentage of the wound interface.

**Figure 5.**
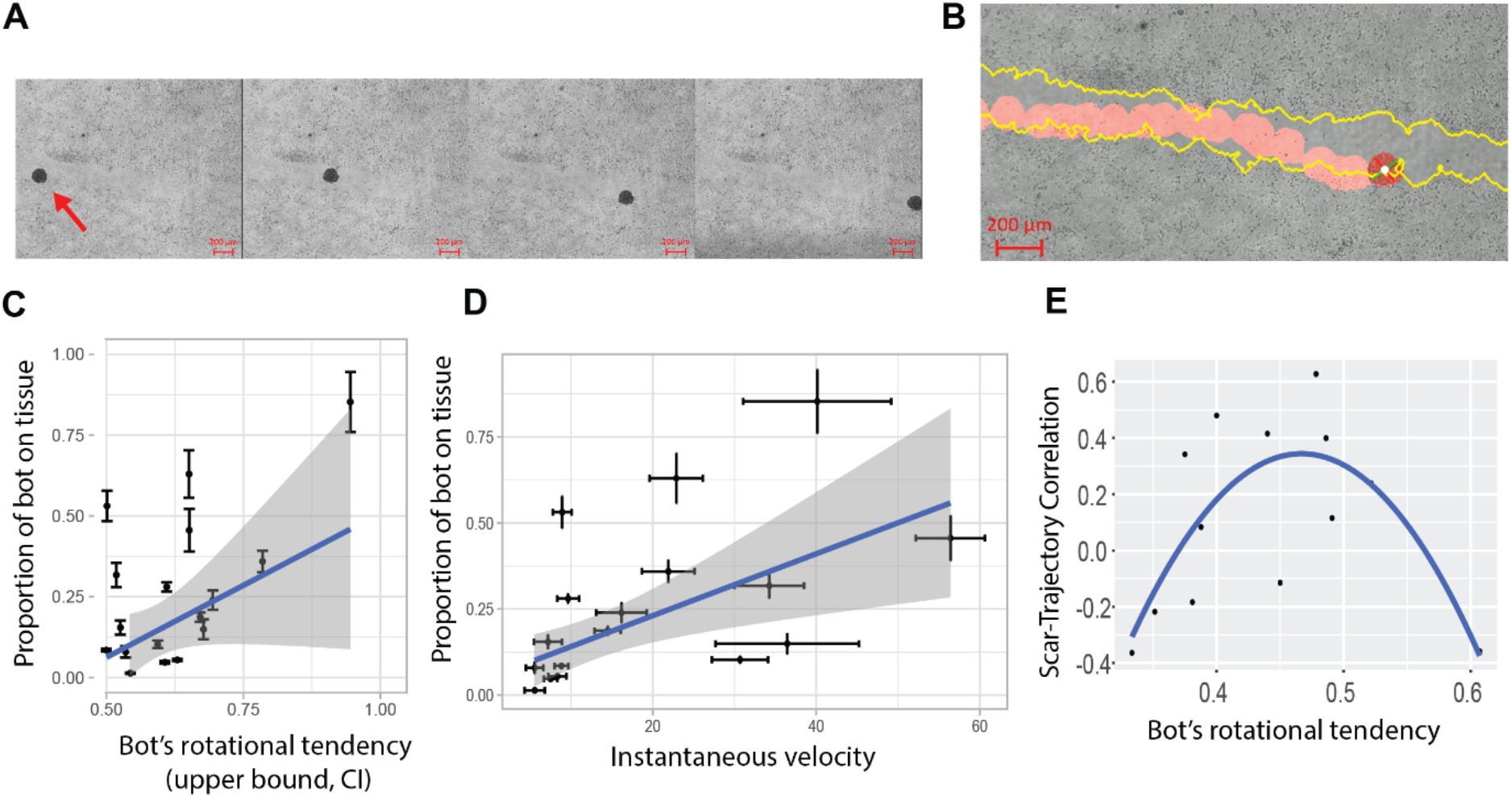
Anthrobots can traverse living tissues in vitro. (A) A representative timelapse video of an Anthrobot as it traverses along a neural wound in vitro. (B) Sample tracking video output with wound edge highlighted in yellow and bot path in red. The rotation of the bot is measured through the change in the orientation of the green and red bars attached to the center of the bot in white. (C) The significant (p=0.017, slope 1.15) positive relationship between bot gyration index and proportion of bot’s body in contact with wound suggest that circular bots explore the edges of the wound more as they traverse along the wound. (D) The significant (p=0.031, slope 0.0082) positive relationship between bot speed and proportion of bot’s body in contact with wound further suggest that faster bots also explore the edges of the wound more as they traverse along the wound. (E) For a subset of bots (dataset constrained to non-stalling bots with rotational tendencies between 0.33 and 0.7 and viable tracking videos), the quadratic relationship (p=0.006) between bot gyration index and wound-trajectory similarity metric suggests that there is a goldilocks zone for how circular bots need to for maximum wound area exploration.

Having observed the behavior of these bots in wounds, we focused on the interactions between various wound surfaces and the bots following the edge of the wound. In order to do so, we first constrained the dataset to trajectories that could be used to further understand the relationships between the two. We specifically focused on bots that were not extreme in their rotational tendencies, had ample contact with the wound and had viable tracking videos (see Materials and Methods). This enabled us to isolate the tendency of the bot to turn consistently in the same direction (measured again by “bot’s rotational tendency”) and the correlation between the trajectory and wound edge (measured by “wound-trajectory correlation”). With our constrained dataset, we saw that Gyration had a quadratic effect on wound-trajectory similarity (Figure 5E) with p=0.006, suggesting that there is a specific range for gyration where the wound-trajectory correlation can be maximized: Anthrobots can be chosen to specifically maximize exploration efficiency based on their gyration.

### Anthrobots can promote regeneration in live tissue wounds

One of the most important aspects of exploring synthetic morphogenesis is the opportunity to observe novel behaviors that are obscured by standard, default phenotypes. Having seen that these airway cell-derived constructs can traverse along and settle in neural wounds, we decided to check for the effects of their presence on the surrounding cells. A characterization of their wildtype capabilities is important not only for understanding biological plasticity but also for establishing a baseline for future efforts in which biobots are augmented with additional synthetic circuits for pro-regenerative applications.

Inspired by collective behavior and swarm intelligence, and more specifically how in nature superorganisms can accomplish tasks that an individual organism cannot, we decided to create “superbot” assemblies by facilitating random self-aggregation of distinct Anthrobots that fuse to form larger structures. We accomplished this without using molds or any other external shape-giving equipment, but by simply constraining multiple Anthrobots in a relatively smaller dish, while keeping everything else constant. Akin to how ants cross openings that are too wide for a single ant to cross by forming a bridge through aggregation of their bodies,^63^ we placed these superbots into arbitrary sites along the tissue wound such that they span the entire width of the wound, enabling them to “bridge” two sides of the damaged tissue in order to see if we can induce any kind of regeneration of the wounded tissue by bridging the two sides, akin to a mechanical stitch. **Figure** 6A shows a superbot on a wounded live tissue upon its placement on day 0, as well as the resulting bridge configuration on subsequent days of day 1 and day 2.

**Figure 6.**
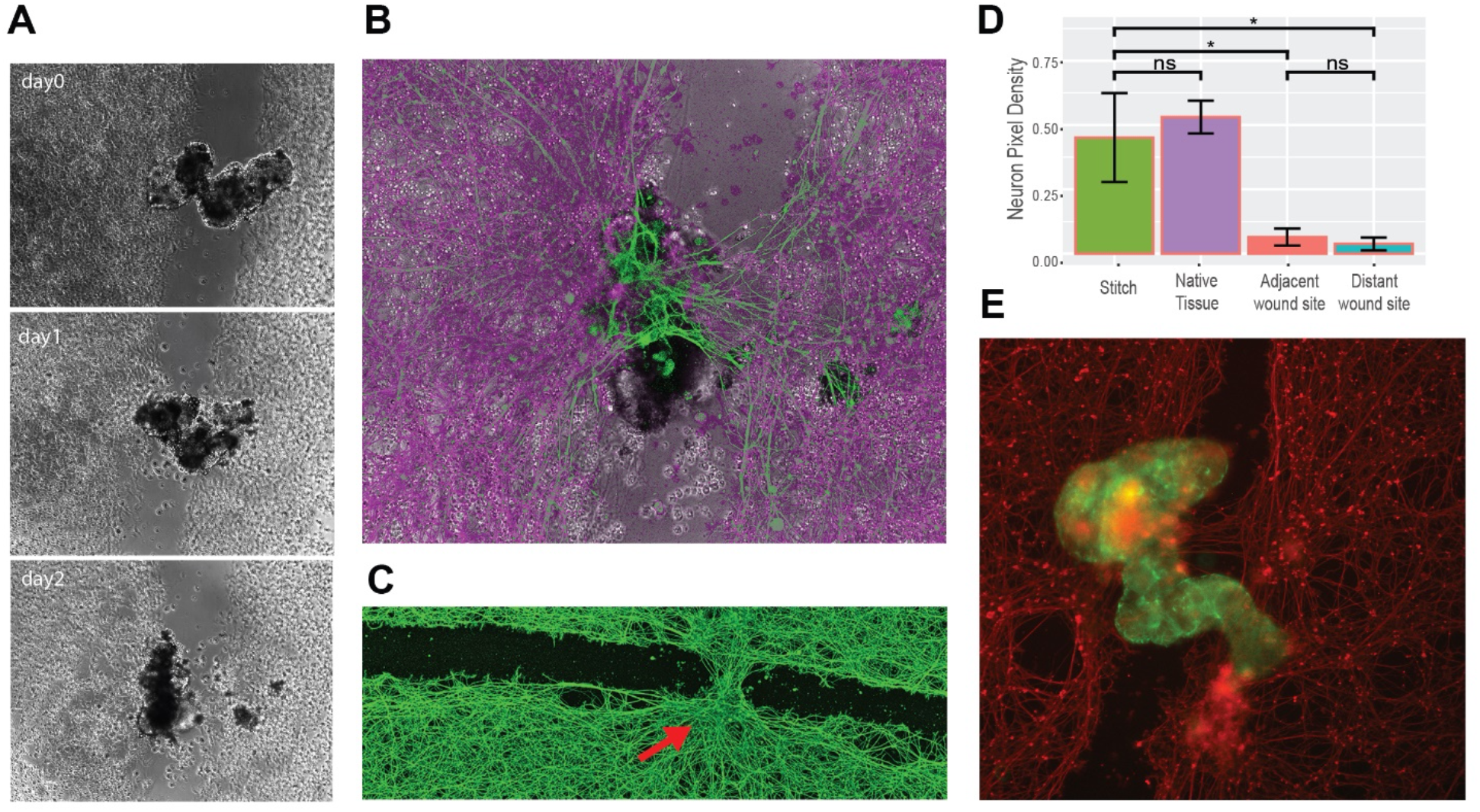
Anthrobots can promote regeneration in live tissue wounds. (A) sample micrograph of a bridge across a neural wound over time. (B) An overlay of a bridge bot and the induced stitch at the end of the observation. (C) Immunological staining of neurons fixed on day3 after the bots were introduced to the system, showing an induced neural repair at the site of bot settlement. Scalebar 500uM. (D) Average proportion of neuronal coverage by pixel counts for each positional category at the healing (self-stitch) site, unwounded native tissue (calculated by the average of the two neuron heavy area pixel coverage), adjacent areas of wound and further reaches of wound. Difference between stitch site and native tissue is insignificant (p = 0.37), while the difference between the stitch site and both adjacent and distal wound sites are significant (w/ p = 0.006 and p = 0.005, respectively) which suggests the healed tissue is as dense as the native tissue, and the healing effect follow a crisp profile as opposed to a gradient profile. See methods section for example frame of a sampling region. (E) Immunological staining of another sample bridge superbot (green) and the neuronal tissue (red).

Strikingly, within the next 72 hours upon inoculation of the superbot into the tissue wound on day 0, we observed a substantial regrowth of the native tissue taking place, resulting in the formation of a stitch right underneath the “superbot bridge,” connecting the two sides of the wound (Figure 6B). This stitch was observed solely at the site of superbot inoculation, and at no other place along the long wound (Figure 6C). A quantitative analysis (Figure 6D) of these self-stitch sites shows that while the neuron pixel coverage density of the stitch is as high as the native tissue outside the wound (see **Supplemental Figure** 4), the rest of the wound space, whether adjacent or far, had significantly less density of coverage. Thus, the density of the induced stitch area that was built with the aid of Anthrobots represented full (statistically indistinguishable from 100%) recovery of the original tissue, and was uniquely different from the surrounding “wound” area: Anthrobot assemblies induce efficient healing of live neural tissue.

## Discussion

Biorobotics and bioengineering have at least two main areas of impact. One is the production of useful living machines.^2–4^ The other is the use of unconventional configurations for living materials at all scales, to probe the macro-scale rules of self-assembly of form and function.^15,64–66^ Specifically, by confronting evolved systems with novel contexts, we can learn about the degree of plasticity that cells and control pathways can exhibit toward new anatomical and functional endpoints, as well as develop protocols to alter default outcomes. Here, we used human patient cells to begin the journey toward immunologically-acceptable, active, living biomedical constructs, and to begin to probe the morphological and functional capabilities of mammalian, adult cells.

Self-motile, fully-organic biobots have been demonstrated with frog cells;^32–34^ however, it was unknown whether their properties depend strongly on their amphibian genome and evolutionary history, as well as their embryonic state. Moreover, their construction depended on a rate-limiting process of extracting source cells from frog embryos. Here we show a protocol for enabling the self-construction of Anthrobots: living structures made from epithelial cells that traverse aqueous environments. The process is highly scalable, and produces Anthrobots in the course of 3 weeks, with minimal manual input beyond weekly media changes. At the end of their 4-6 weeks life span, they safely degrade.

Anthrobots exhibit several distinct movement and morphological classes, which are significantly correlated. This is especially important because the structure and function of this novel construct is not that of a familiar organism (despite a wild-type genome), and it was not yet known whether its morphospace possessed specific attractors, how reliable the cells’ navigation of that morphospace was, or how the movement patterns would relate to its specific morphology. Such correlation also has implications for future control of higher-order behaviors (such as movement types) by way of controlling Anthrobot morphology through synthetic morphogenesis, as well as real-time physiological signaling. Analysis of movement and morphology revealed the ability of the Anthrobots to establish bilateral symmetry, which is an interesting aspect of self-assembly in a symmetrical environment, and will enable future studies of the still poorly-understood question of how multicellular amniote embryos bisect themselves to establish a single midplane for their bodyplan.

Anthrobots are derived from adult human tissue, and in the future could be personalized for each patient, enabling safe in-vivo deployment of these robots in the human body without triggering an immune response. Once inoculated in the body via minimally invasive methods such as injection, various applications can be imagined, including but not limited to clearing plaque buildup in the arteries of atherosclerosis patients, bulldozing the excess mucus from the airways of cystic fibrosis patients, and locally delivering drugs of interest in target tissues. They could also be used as avatars for personalized drug screening^67^ having the advantage of behavior over simple organoids, which could be used to screen for a wider range of phenotypes.

## Conclusion

These data establish a research program with many unanswered questions for subsequent work. What other cells can Anthrobots be made of? What other behaviors might they exhibit, and in what environments? What other tossie types can they repair or affect in other ways? Can transcriptional or physiological signatures be read out in living bots, that reflect their past and immediate interactions with surrounding cellular or molecular landscapes? Do they have preferences or primitive learning capacities,^68^ with respect to their traversal of richer environments? More fundamentally, these data reveal additional morphogenetic competencies of cells which could have implications for evolutionary developmental biology, as evolution of anatomical and functional features could be affected by the ability of the same genome to produce very diverse forms in different environments. Finally, this kind of new model system is a contribution to two key future efforts. The study of synthetic biological systems^9,15,69,70^ is an essential complement to the standard set of phenotypic defaults available in the natural phylogenetic tree of Earth, revealing the adjacent possible in morphological and behavioral spaces.^71,72^ Moreover, these systems offer a safe, highly tractable sandbox in which to learn to predict and control the surprising and multi-faceted system-level properties of multiscale complex systems.

## Materials and Methods

### NHBE culture

Normal human bronchial epithelial cells (NHBE) were used from Lonza Walkersville, MD. The cells were first thawed and seeded on a T150 flask containing bronchial epithelial growth medium (Lonza CC-3170) for 2D cell culture growth. Once the NHBEs were ~80% confluent, they were passaged into a 24-well-plate of Matrigel beds for 3D cell culture. The NHBEs were not passaged past the 3^rd^ passage. Each matrigel bed contained 500μL of 25% Matrigel-, 0.1% 0.5nM retinoic acid and 75% BEDM that was centrifuged for 5 seconds at 100 × g and prepared at least 4 hours before passaging the cells. The cells were re-suspended in 5% matrigel, 95% BEDM and 0.1% RA and seeded directly onto the matrigel beds with 500μL per well at a 30,000 cells/mL concentration. Once seeded, the NHBEs were centrifuged for 5 seconds at 50 x g. On days 2 and 8, the NHBEs received a top feed containing 750μL 5% matrigel, 95% BEDM, and 0.1% RA. On day 14, 500μL of the wells’ contents was aspirated and 500μL of dispase (#D469) at concentration 2 mg/mL was added to each well. A mini cell scraper was used to break up the matrigel clumps and then followed by a 0.05% Triton coated pippete tip to mix up the matrigel with the dispase. The dispase was then incubated at 37°C for 1 hour with the pipetting process repeated every 15 minutes. During incubation, Pluristrainer Mini’s with a 40 μm pore size (Fisher Scientific #431004050) were placed in wells of a fresh 24-well-plate that contained 2.5 mL of 0.05% Triton. After the incubation period, 250 μL of 1%5mM EDTA in D-PBS was added into each well. The media in each well was then drawn up, using the Triton-coated tip, and added to the Triton-coated strainers. The NHBE spheroids in the strainer were rinsed with 1 mL of D-PBS then expelled onto a low adhesive dish by inverting the strainer over the dish and expelling 5 times of 1 mL of BEDM through the bottom of the strainer. See the “Anthrobot Maintenance in Culture” section of the Supplemental Methods Information for more details.

### Immunocytochemistry / Immunofluorescence

Anthrobots were collected in Pluristrainer Mini’s with a 40-micron pore size (Fisher Scientific #431004050) and fixed with 4% paraformaldehyde at room temperature for 30 minutes. Immunological staining was carried out with standard methods using 10% normal goat serum with 1% BSA, as well as the following antibodies mouse anti-acetylated tubulin (Sigma-Aldrich #T7451) Alexa Fluor 594-conjugated mouse anti-ZO-1 (Thermo Fisher Scientific #339194). Images were collected using a Lecia SP8 FLIM microscope with a 25× water immersion objective. Z-stack step size=3μ unless otherwise specified. See the “Anthrobot Maintenance in Culture” section of the Supplemental Methods Information for more details.

### Tracking timelapse videos

Timelapse videos of the Anthrobots were contrast-enhanced using the video editing software ImageJ. They were then processed to extract the trajectories of the Anthrobots utilizing the trackR function in the trackR package (version 0.5.1) for R. The software parameters were chosen manually in order to increase the accuracy of the tracking, following the instructions in the trackR package’s help. Tracking errors such as the swapping, deletion or insertion of tracks were subsequently manually corrected using the trackFixer function from the same package.

### Movement type analysis

From the extracted trajectories, we computed the following metrics: (i) the linear distance between the current position and the immediately preceding one; (ii) the linear speed at each position, approximated as the distance moved between the current position and the immediately preceding one during the time interval between these two positions; (iii) the heading of the bot at each position, approximated as the angle between the vector formed by the current position and the immediately preceding one and that formed by the Anthrobot position and the immediately following one; (iv) the angular speed of the bot at each position, approximated as the difference between the heading at the immediately preceding position and that at the current one during the time interval between the corresponding three positions required to calculate these two headings; (v) the time difference between each position. See the “Movement Types Analysis” section of the Supplemental Methods Information for more details.

### Morphotype analysis

To find whether there were any unique morphological types like was the case with movement types, each spheroid was processed through a custom-made analysis pipeline (code attached). First, we created a 3D model of the bot in R where the points corresponding to the body and the cilia were clearly indicated. For this, we first isolated the cilia channel from the LIF images of the bots and ran the images through CiliaQ^73^ using the RenyiEntropy algorithm for detection. These binarized cilia were then imported into our code, along with the points that comprised the body.

See the “Calculation Methods for Morphotype Indices” section of the Supplemental Methods Information for more details.

### Morphotype & movement type correlation analysis

After finding trends among behavioral and morphological data, we decided to see if there was any potential overlap between the two and performed a Fisher test to compute correlation between the major morphological and behavioral classes. See the “Fisher Test for Morphology and Behavior Correlation Analysis” section of the Supplemental Methods Information for more details.

### Bilateral Symmetry Along Movement and Other Axes

After realizing that polarity could play a key role in determining movement type, we decided to see whether the symmetry across the movement axis had any trends compared to the other axes. To calculate this, we followed the procedure used in both Fig 3 and Fig 4 to get representations of Cilia on the body of the bot, then project these representations onto the plane of symmetry defined by the points obtained in the Motility orientation alignment section. The side of the plane (movement axis) each cilia point belonged to was noted using the sign of the dot product of the normal of the plane and the vector to the cilia point. Finally, to better distinguish whether the cilia distribution played a role in movement type (linear vs. circular) we created the Bilateral Symmetry index. See the “Bilateral Symmetry” section of the Supplemental Methods Information for more details.

### Neuronal Traversal and Regeneration

To better explain the relationship between bot trajectory and movement in certain environments, we tried to relate the wound edge to the actual movement of the bot. The steps taken before analysis involved (i) creating a background image prototype from the.czi recording, (ii) verifying the quality of the background image and saving as .png file, (iii) tracking the bot, (iv) extracting the coordinates of the wound, and (v) checking whether tracking was correctly carried out by generating a video with a beacon on the bot. See the “Neuronal Traversal and Healing Tissue Density Analysis” section of the Supplemental Methods Information for more details.

## Supporting Information

Supporting Information is available from the Wiley Online Library or from the author.

## Acknowledgements

We thank Santosh Manicka for helpful technical discussion of Multivariate Classification, Roger Kamm and David Kaplan for in-vitro bot inoculation discussion. We also thank Doug Blackiston, Joshua Bongard, Caitlin Grasso and Sam Kriegman for high-level scientific discussions, Cindy Zhu for image processing help, and Julia Poirier for helpful comments on the manuscript.

## Author contributions

Experimental design and data interpretation: GG, ML. Data acquisition and analysis: GG, PS, BC, HL, BS, SG. Writing manuscript: all co-authors. Funding Acquisition: ML.

## Supplemental Text

### F2-Metrics Breakdown for Different Movement Types

As a result of these behavioral characterization analyses, we observed that the circular bots (type 1, Figure 2D) score the highest on gyration and lowest on straightness indices (Figure 2F). They also have highly similar trajectories and are very common among the behaviors of Anthrobots given this cluster has the smallest homogeneity and a representation of over 30% of all the recorded periods (Figure 2E). We also observed that the linear bots (type2, Figure 2D) score the highest on straightness and lowest on gyration indices (Figure 2F). They have less homogeneity than circular bots but also have the greatest representation out of all clusters (Figure 2E). Accordingly, circular and linear bots together make up more than half of the population, and each have highly homogeneous populations. Finally, the third most common (Figure 2E) type of bot is the curvilinear bot (type 3, Figure 2D), which scores high on both the gyration and straightness indices (Figure 2F) and has the second most heterogeneous trajectories (Figure 2E). Bots with most disorganized trajectories and smallest representation in the overall population (Figure 2E) are the eclectic bots (type 4, Figure 2D), which score the lowest on both the gyration and straightness indices (Figure 2F) due to exhibiting eccentric trajectories that are often a combination of the other three types.

### F2-PanelG: Markov Chain Model

Markov chain model shows state transitions between different clusters and the degree of commitment to a given behavior (persistence), with the circular bots (type 1) as the most committed category with 92.1% chance of the next period being a circular if the current period is a circular. It is followed by linears and curvilinears, which are also relatively consistent at 80.0% and 75.3% respectively. Cluster 4, or the eclectics, as expected, are very unstable, with a consistency of only 39.6%. Cluster 4 seems to act as a sort of intermediate, since there is a substantial chance of the eclectics converting to linear (34.5%) or to a lesser degree circular (15.0%) or curvilinear (10.7%). The transition probability between circulars and linears and vice versa is the lowest and almost nonexistent, at 0.3% and 0.2% respectively. Linears, curvilinears, and circulars rarely convert into eclectics with a probability of 12.3%, 7.5%, 5.8% respectively (and when they do, it is most likely due to collisions or using eclectics as an intermediary).

## Supplemental Figures

**Figure S1.**
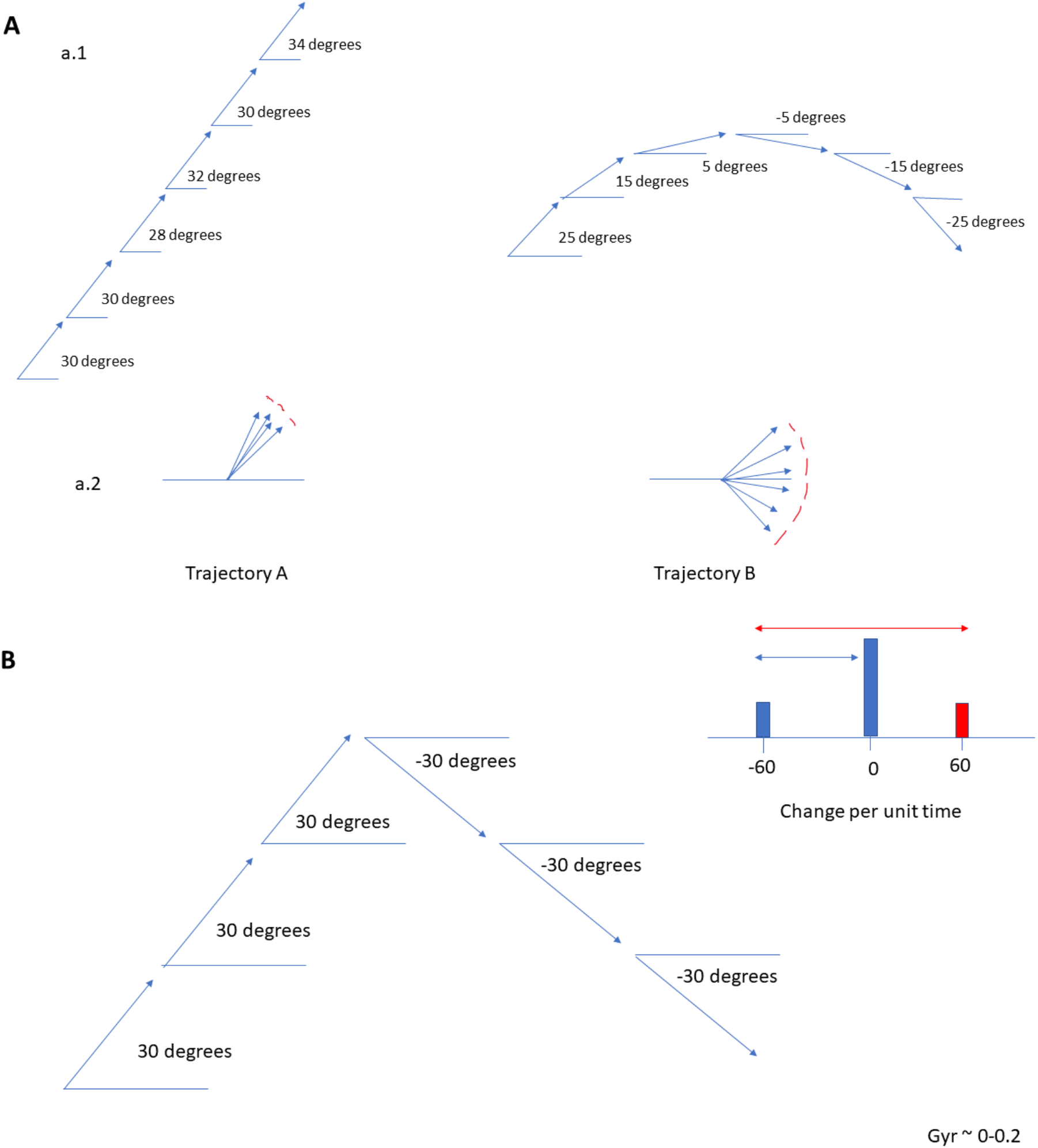
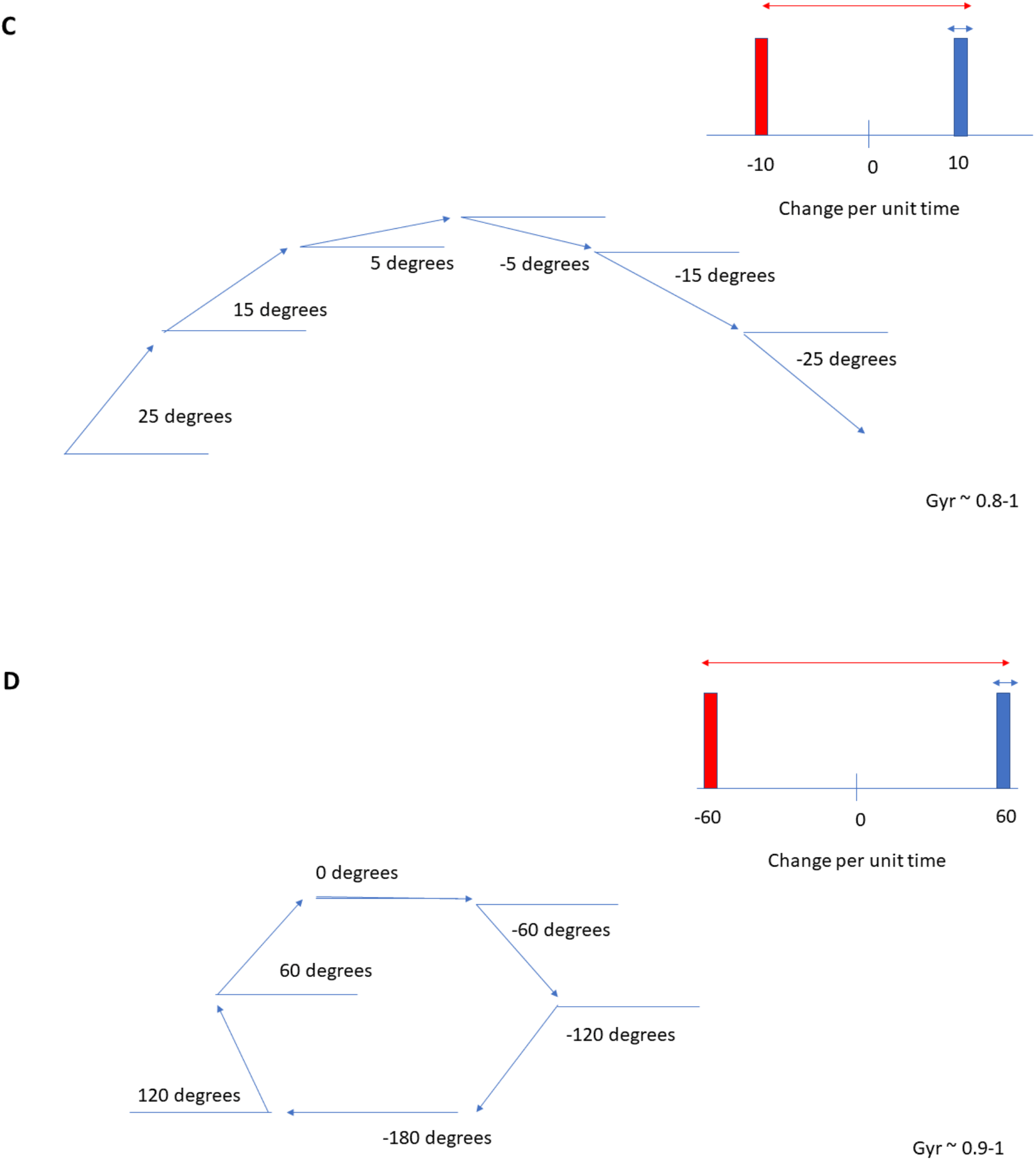
Sample trajectories that reference a relative x-axis to find heading angle. (A) Visual representation of straightness index, which calculates spread of headings as a whole without taking into account temporal dynamics. (B, C, D) Visual representation of gyration index, which includes the temporal aspect and calculates spread of change in headings relative to their magnitude. Graphically, it is represented as the ratio of the blue length (original circular variance) to the red length (circular variance of the original angular speeds and their additive inverse) on the histograms.

**Figure S2.**
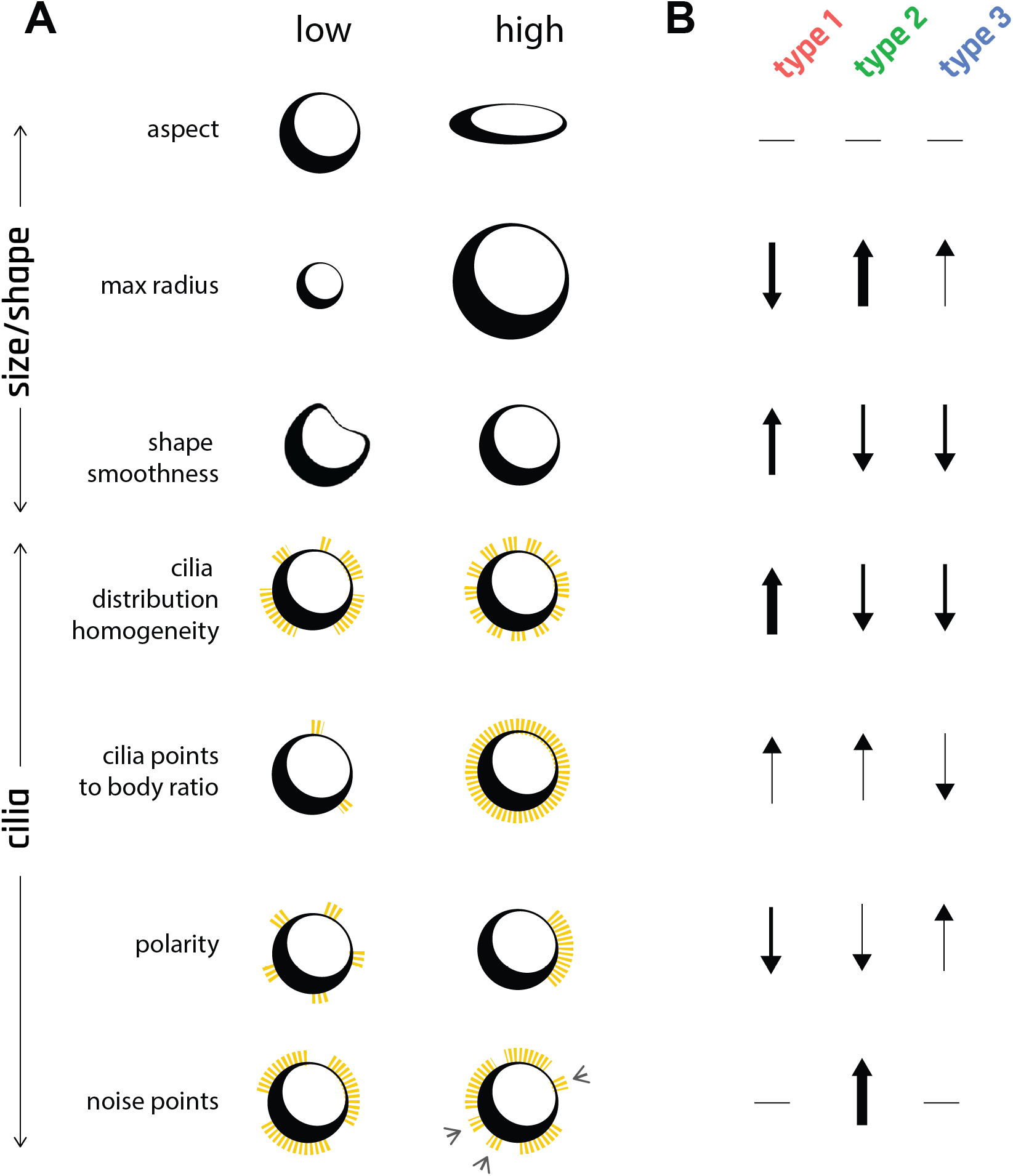
Eight morphological indices were used to characterize Anthrobot morphotypes. (A) Visual summary of morphological indices at their extreme points (low, high). (B) Graphic summary of boxplots on Figure 3D, describing 3 different morphotypes. Arrow thicknesses are correlated with the # of standard deviations between a given morphotype’s mean vs the overall population mean for a particular morphological index.

**Figure S3.**
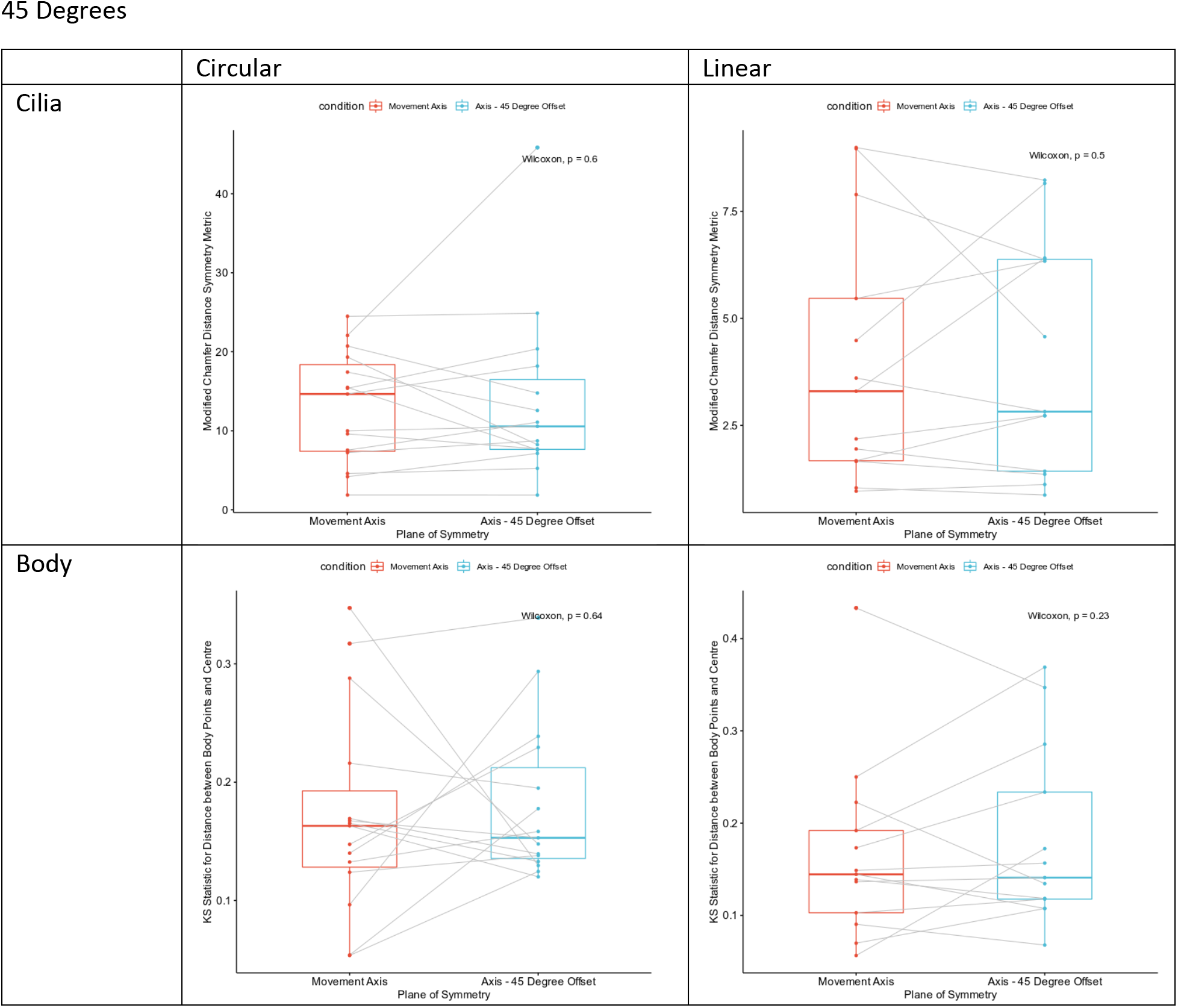
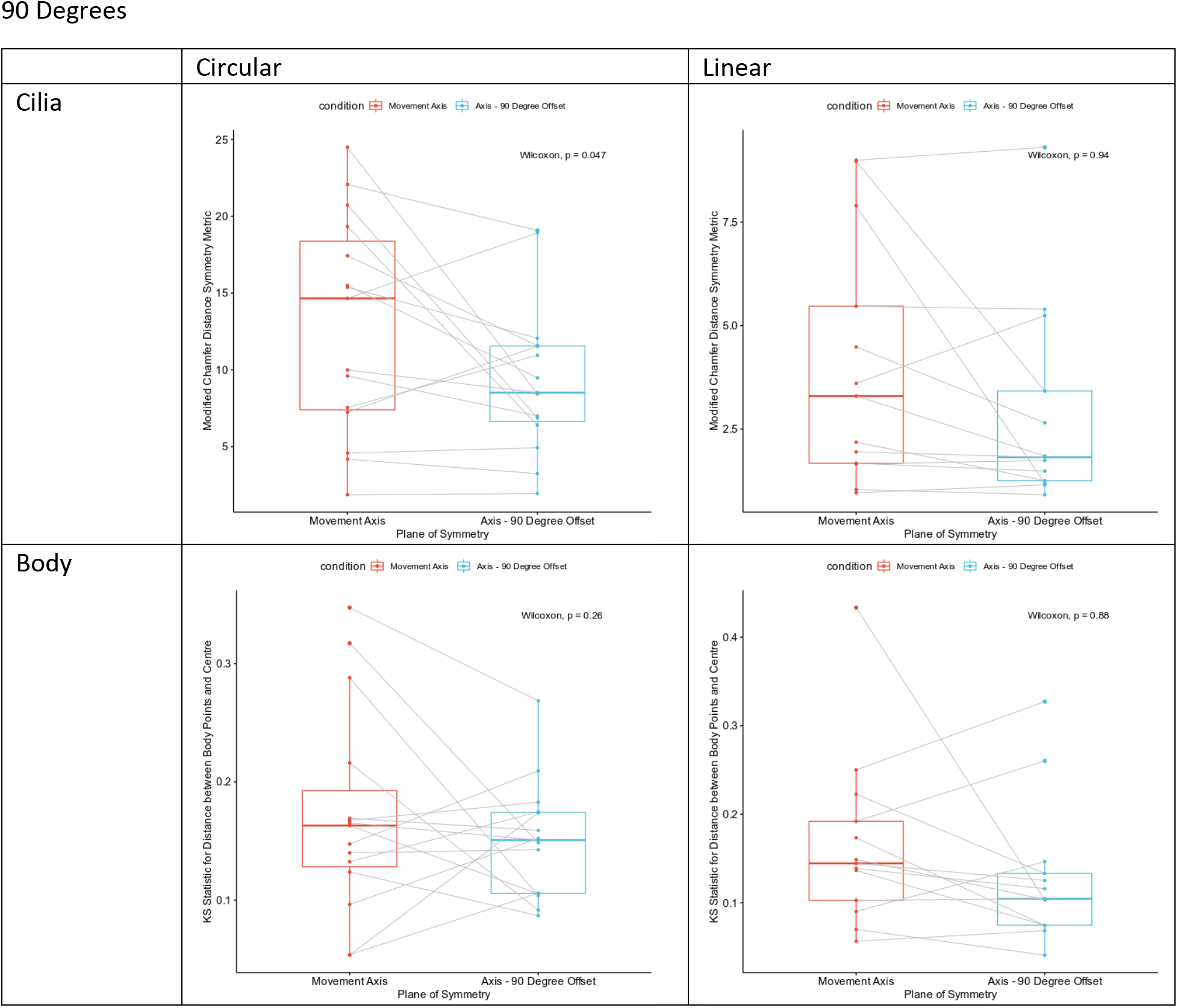
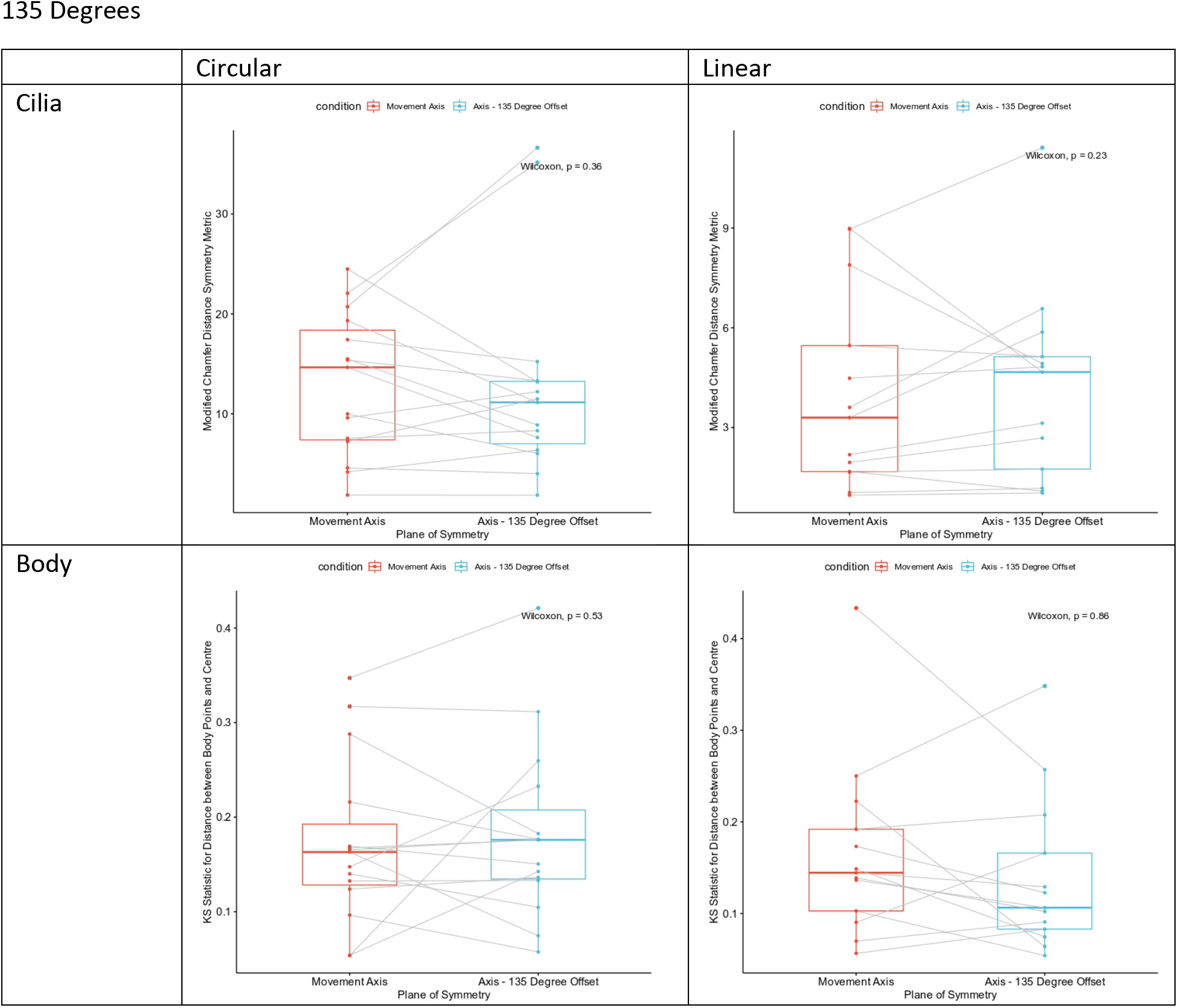
Difference in asymmetry of cilia distribution and body shape between linear and circular bots in respect to the movement axis and its 45, 90, and 135-degree offset axes.

**Figure S4.**
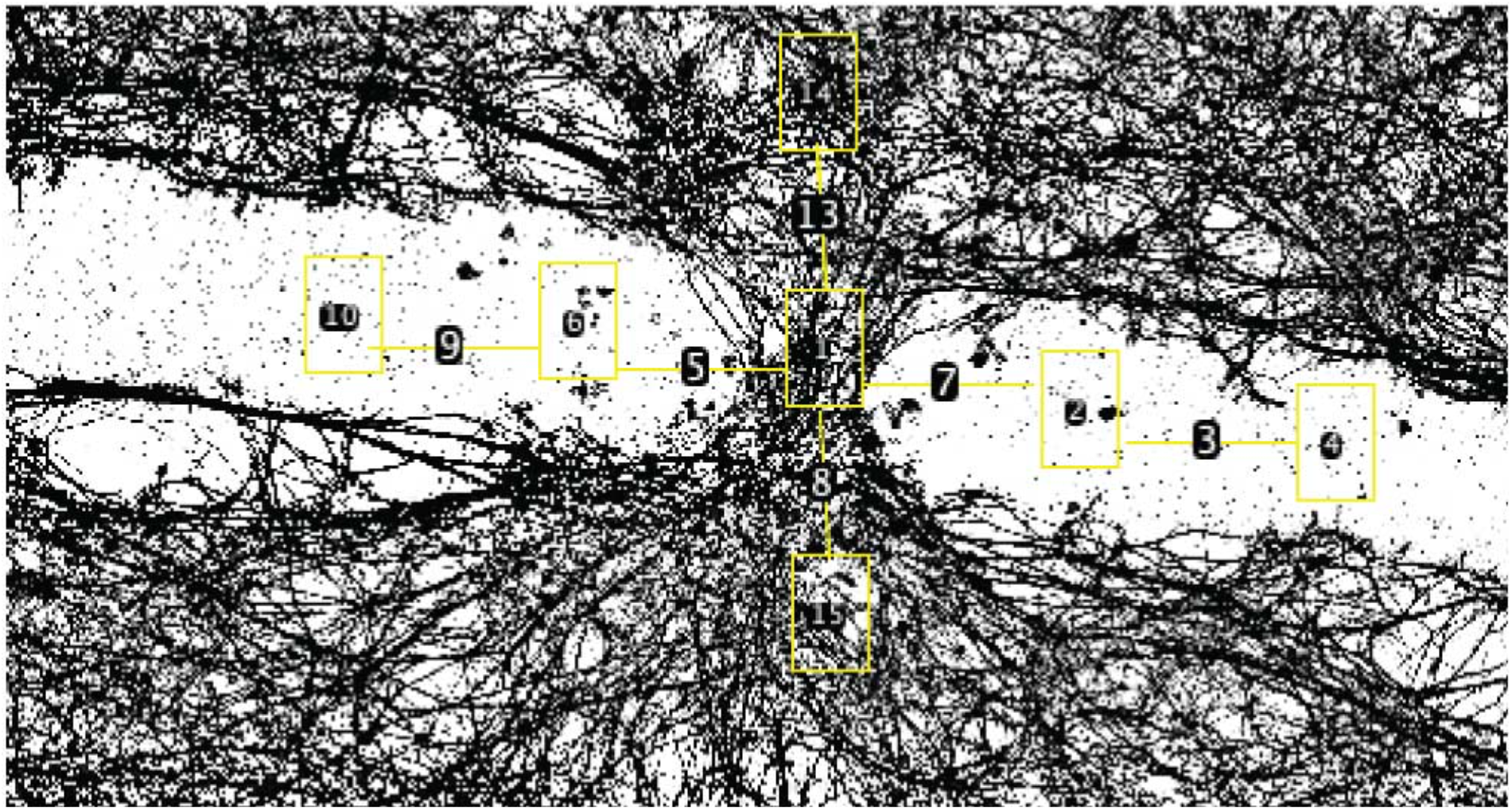
Sample neuronal density sampling region. Each rectangle represents an area sampled and the lines are consistently the same length, the “bridge length.”

## Supplemental Methods Information

### Anthrobot maintenance in Culture

After all spheroids were in one dish, they were divided evenly amongst multiple 60mm dishes by using a Triton-coated pipette tip and a microscope to manually draw up and divide them. 0.5μL of 0.5nM retinoic acid was added into each dish once divided. For the next 14 days as the spheroids started moving, they required 0.5μL of 0.5nM retinoic acid every other day and a media change every 4 days. The media change was performed by circling the Anthrobots to the center of the dish then collecting 2 mL of old media and adding 3 mL of fresh BEDM-this was done under a microscope to ensure no Anthrobots got aspirated.

### Immunocytochemistry / Immunofluorescence

Anthrobots were collected in Pluristrainer Mini’s with a 40-micron pore size (Fisher Scientific #431004050) and fixed with 4% paraformaldehyde at room temperature for 30 minutes. Following PBS washes, blocking and permeabilization were performed for 1 hour at room temperature on a rocker in a blocking buffer consisting of phosphate-buffered saline with 10% normal goat serum, 1% bovine serum albumin, and .15% triton x-100. Anthrobots were then incubated with mouse anti-acetylated tubulin (Sigma-Aldrich #T7451) and rabbit anti-cytokeratin 5 (Abcam #ab52635) primary antibodies diluted 1:250 and 1:100, respectively, in blocking buffer for 24 hours at 4oC on a rocker. The primary antibodies were labeled with Alexa Fluor 647 donkey anti-mouse (Thermo Fisher Scientific #A31571) and Alexa Fluor 488 donkey anti-rabbit (Abcam #ab150073) secondary antibodies, both at 1:500 dilutions in blocking buffer, for 1 hour at room temperature on a rocker. Lastly, Anthrobots were incubated with Alexa Fluor 594-conjugated mouse anti-ZO-1 (Thermo Fisher Scientific #339194) at a 1:100 dilution in blocking buffer for 24 hours at 4oC on a rocker. Anthrobots were mounted on glass-bottom 96-well plates in ProLong Glass Antifade Mountant with NucBlue (Thermo Fisher Scientific #P36981). Images were collected using a Lecia SP8 FLIM microscope with a 25x water immersion objective. Z-stack step size=3 micron unless otherwise specified.

### Neuronal Culture

Followed a previously established method(*1*). A 150 cm dish was first coated with 0.1% gelatin for 20 minutes and then aspirated off before seeding mouse embryonic fibroblasts (ATCC #SCRC-1008) in MEF growth media (89% DMEM GlutaMAX, 10% Fetal Bovine Serum, and 1% Anti-anti). Once the MEFs were confluent, they were inactivated by adding 20 mL of MEF growth media containing 500 μL of 10μg/mL mitomycin C (Sigma #M4287) and incubating for 2-3 hours at 37°C. After incubation, the MEF growth media + mitomycin C media was replaced with hiNSCs at a density of 1/10 of a confluent target vessel in 25 mL of hiNSC growth media (77.6% Knockout DMEM, 20.20% KOSR, 1% GlutaMAX, 1% Anti-anti, 0.18% 2-mercaptoethanol with 0.1 % of 20 ng/mL bFGF). The day after seeding the hiNSCs required a media change where all the old media was aspirated off, and 25 mL of fresh hiNSCs growth media was added. Media changes were performed every other day until the hiNSCs were 80-85% confluent. 3 hours before performing the differentiation, the destination vessels were first coated with .1mg/mL poly-d-lysine (enough to coat the bottom of the wells) for one hour at room temp, and then the PDL was aspirated before adding in 10ug/mL laminin in DPBS (enough to coat the bottom) for 2 hours at 37°C. In the differentiation, the hiNSCs first went through one D-PBS wash before adding TrypLE Select for 3-5 min to detach the cells from the plate. The cells were then collected and spun down for 3 minutes at 500g then resuspended in neurobasal differentiation media (96% Neurobasal Media, 2% B-27 supplement, 1% GlutaMAX, 1% Anti-anti). The hiNSCs were seeded at a concentration of 100,000 cells/cm^2. Once the hiNSCs were in differentiation, there was a media change the day preceding their differentiation and then every other day from there.

### Movement types Analysis

Behavioral classification was performed on non-overlapping 30-second blocks of trajectory. To determine how predictable a position change was, the linear speed, heading, and angular speed were estimated at each position to predict the coordinates of the following position. The error (Euclidean distance) between the predicted coordinates and the actual coordinates was then computed. For each complete 30-second block of trajectory (i.e., a block with no missing timestamp), total error over the entire block was calculated and normalized by the total distance traveled during that block to account for the artificial error amplification caused by predicting over longer distances. To separate active from inactive blocks, an automated classification method was used on the distribution of total normalized errors. A gamma mixture model with two components was fit to the data using the expectation maximization algorithm in the REBMIX function from the rebmix package (version 2.12.0) for R.^1^ The 30-second periods in the resulting cluster with the highest total normalized error were considered as inactive and excluded from further classification.

We then derived two metrics to describe each trajectory: (i) a “straightness” index computed as 1 minus the circular variance of the headings during the block (a value of 1 indicates a perfectly straight line) and (ii) a “gyration” index computed as 1 minus the circular variance of the angular speed during the block divided by the circular variance of the same angular speeds and their additive inverse, which helps in taking into account the magnitude of the angular speeds themselves (a value of 1 indicates a trajectory following a perfect circle).

The need for two indices arises from the fact that a straightness index alone cannot fully tease apart all different movement types due to its aggregate view of a trajectory. In other words, a low straightness index does not automatically translate into a perfectly circular bot, as we can see in Supplemental Figure 1-a.1 and 1-a.2 with the arc trajectory. This is due to the fact that the straightness index does not account for the time-dependent dynamics and thus ignores individual variations across frames. This is where the second movement metric, the gyration index, comes into play. To account for temporal information, we calculate the angular speed, which is the difference between successive headings divided by time between frames and thus has units of radians/ second. Supplemental Figure 1B shows a bent trajectory and Supplemental Figure 1C shows an arc trajectory, both of which have similar straightness indices. However, when we start looking at their temporal relationships using angular speeds, the behavior is entirely different. For the arc (Figure S1C), the variance of the angular speed is very small since the change in heading of the trajectory each time is relatively consistent (the distribution shown in the histograms).

For the bent trajectory (Figure S1B), the variance of angular speed is much larger than the arc since for most of the trajectory the angular speed is close to 0 (it goes straight), but the bent portion has a very high angular speed, i.e., the angle changes very quickly. In general, the greater the absolute value of the angular speed the sharper the turn in the trajectory (zero is straight) and the greater the variance of the angular speeds, the less the consistency of the turns in the trajectory. A circle or arc usually has absolute values of the angular speed much greater than zero and low variation of angular speed. However, the gyration index alone cannot differentiate between all behavior either. Let’s look at a circular trajectory (Supplemental Figure 1D). Even though the absolute values of the angular speeds between arcs and circles are different, the circle also ends up having a gyration index close to 1 since all the turns in a circle are highly consistent like in an arc and thus the variance of the angular speed for both is very small. This fact means the gyration index cannot segregate between arcs and circles, among other things, by itself. Interestingly, the straightness index is exceptional at separating arcs and circles. This shows that though either index alone cannot distinguish all movement types well, together they can accomplish much more.

To separate the trajectory blocks into categories of similar behavior after calculating the movement metrics, a cross-entropy clustering algorithm was used,^2^ and implemented in the cec function of the CEC package (version 0.10.2) for R.^3^ This yielded us six categories, of which trajectories from categories numbered 3 and 4 were merged into categories numbered 1 and 2 respectively due to the difference being phenotypically minimal. In the “behavioral space” as defined by the straightness and gyration indices, cluster 3 had the same straightness index range as cluster 1 and a lower gyration range between roughly 0.65 and 0.95, which represented trajectories that were highly circular but fell short of cluster 1 which represented “prototypical circulars”. Similarly, Cluster 4 had a slightly smaller straightness index range than cluster 2 (0.7 to 1 instead of 0.6 to 1) and higher gyration range between 0.1 and 0.55 which represented trajectories that were mostly linear but didn’t have a high enough gyration to be curvilinear or low enough gyration to be Cluster 2, a “prototypical linear”. The merge of the two clusters increased the average dissimilarity of the cluster, but it is a testament to how similar clusters 1 and 3, and 2 and 4 were already that their dissimilarity still remains very low at ~0.09 and ~0.14 respectively. Last, to understand how the bots’ behaviors are distributed relative to each other, transition probabilities between each behavioral category were estimated by calculating the proportion of times a block of a given category is followed by a block of the same or another category. This was then presented in the form of a Markov Chain.

### Calculation Methods for Morphotype Indices

The “body points’’ were extracted using the “body channel” of the LIF images by first running the pixels through a logistic transform then thresholding the pixels based on the signal to noise ratio, calculated by using a median filter and comparing the points before (“signal + noise”) to after (“only signal”).

Due to the large volume of body points, to reduce the points to a manageable amount, we found outlines of each slice of the spheroid by using a concave hull using the Concaveman package (version 1.1.0).^4^ The cilia points were then projected on the nearest body points by simply choosing the nearest one by distance to get the shadow of the cilia on the body.

The structural index variable Cilia Points was calculated by counting the number of unique projected points on the body. Then, the dbscan package (version 1.1-10)^5^ function in R was used to find the clusters of cilia. The number of points that fell outside of clusters with this definition were defined as Noise Points.

Following this step we computed the spanning ellipsoid of the body points by using the ‘ellipsoidhull’ function from the cluster package (version 2.1.3).^6^ The Max Radius variable was calculated directly by the function, and Aspect was defined as the ratio of the largest radius to the shortest radius, all quantities computed by the function. Finally, we used the ‘ashape3D’ function from the alphashape3d package (version 1.3.1)^7^ to generate a 3D alpha hull of the body points, and used the mesh to get the surface area of each spheroid.

Cilia Points/Area was defined as the Cilia Points variable divided by the calculated surface area. Similarly, the Shape Smoothness was defined as the ratio of the volume of the 3D alpha hull to the volume of the spanning ellipsoid. Finally, we found the center of the bot by finding the sum of the centroids of each triangle that makes up the alpha hull weighted by area of the triangle. Polarity was defined as the norm of the vectors from the center to each cilia point divided by the sum of the norm of each vector. The Cilia Distribution Homogeneity was defined as 1 -D statistic of the two sample Kolmogorov-Smirnov test, where sample A is the 1^st^ nearest neighbor (1NN) distances for the cilia, and sample B the 1NN distances if the same number of cilia points were distributed close to uniformly but randomly across the surface of the bot. After these analyses were carried out, we ran the dataset through a Principal Components Analysis with centering and scaling. Afterwards, a hierarchical clustering was carried out on the resultant dataset with the Ward.D2 method and the resulting classification was plotted as above. In total, 350 bots were put through the pipeline and into the following PCA and included Movers, Nonmovers, Linears and Circulars. Further details can be seen in the code.

To get a confidence interval for the absolute value of the loadings, we bootstrapped the loading value by sampling 250 bots from the 350 that we have, 10 times. We then took the loadings value for the 1^st^ and the 2^nd^ primary component for all 8 variables and calculated the mean and 95% Confidence Interval for the loadings for the PC in question. If there was overlap between the CI of the loadings, they were assigned the same rank, otherwise they were assigned different ranks. Ranks were relative to the “highest” contributor of the rank; i.e, for PC1, Shape Smoothness had overlap with Max Radius, but it also had overlap with Cilia Distribution Homogeneity. However, Cilia Distribution Homogeneity did not overlap with Max Radius. Max Radius had the highest upper limit of the CI among the 3 variables in question, and thus Shape Smoothness was co-ranked #1 along with Max Diameter, but Cilia Distribution Homogeneity was not.

### Fisher Test for Morphology and Behavior Correlation Analysis

After finding trends among behavioral and morphological data, we decided to see if there was any potential overlap between the two. To eke out any possible correlation, we first chose to use categories of spheroids behaviorally orthogonal to movers, the non-movers. The goal was to observe whether there is any overlap between the morphology of movers and non-movers. Similarly, we had four potential behavioral types that could overlap with our morphological clusters. Eclectics could not be included in the analysis since they are an aggregate of multiple inconsistent patterns and highly uncommitted to their behavior (Figure 2G), and thus cannot be used in a bot-level analysis (instead of period). Circulars and Linears on the other hand were highly committed behavioral types that were orthogonal to each other (had little to no interconversion on the Markov plot) and prototypes of two extreme movement types with high variability between them. Consequently, the Curvilinear behavioral subtype was also not included since it lacked orthogonality with both Circulars and Linears due to shared traits between them. The morphological indices for each spheroid were calculated as outlined in the methods for the previous sections, and then clustered with Circulars, Linears, Nonmovers and Movers together. To measure the significance of the overlap, if any, between clusters, we decided to use a Fisher test to compute whether the proportion of a certain behavior per cluster type was different from the others. We ran the test twice, once to see if there were any significant differences in number of nonmovers per cluster, and once to compute the difference in the ratio between circulars and linears per cluster. It showed that the proportion of nonmovers in Cluster 1 vs. Clusters 2 and 3 were significantly different with an average p=2.6*10e-6 and 3.5*10e-8 respectively Cluster 2 and 3 also had a statistically significant difference in number of nonmovers (p=0.01) which is understandable, since Cluster 2 had no non-movers. For Circular/Straight, Cluster 2 vs. 3 were significantly different with p=0.00011, and Cluster 1 had no Circulars nor Straights.

### Bilateral Symmetry

Bilateral Symmetry index was modified from the Chamfer distance, and was calculated as the sum of the median/ mean of the distances between all points in set A and the closest point in set B and the median/ mean of the distances between all points in set B and the closest point in set A. To calculate the index, the cilia points were projected onto the plane defined by the three points in the section below. Set A and Set B then became the points projected from one or the other side, respectively, after which the modified Chamfer index was calculated for the two sets using the createTree() function of the SearchTrees package. The statistics used to calculate the asymmetricality between both sides were the difference in points between the two hemispheres, the difference in points/ total cilia points, the median and the mean modified Chamfer distance. In the end, they were visualized and clustered using a PCA to see trends (see code).

In the case of figure 4E, instead of calculating the asymmetry statistics after getting the equation of the plane, we then used the Rodrigues’ rotation formula to rotate the normal (and thus the plane) with the fixed intersection being the center of the bot. Rotations of 45, 90 and 135 degrees were used yielding 4 axes (in the form of plane equations) including the movement axis. To eliminate the z-axis we used a PCA to get the rotation matrix to convert the projected cilia points from 3D to 2D and calculated the Chamfer distance using the formula described at https://github.com/UM-ARM-Lab/Chamfer-Distance-API with the exception of we didn’t square point distances. This statistic was calculated along for the cilia of all linears and circulars and put into a paired Wilcoxon rank-sum test with an alternative hypothesis of “greater” and “less” for circulars and linears respectively to see if the movement axis was “more asymmetrical” or “less asymmetrical” respectively. For the body we did the same procedure, with the exception that our statistic now involved finding the distance of the body points from the centre of the bot (once again segregated into two hemispheres with the dot product). Then, we used the KS test to calculate a D-statistic which had greater values the more dissimilar the two distance distributions for both hemispheres were. Just like the cilia we then used a paired Wilcoxon rank-sum test with an alternative hypothesis of “greater” and “less” for circulars and linears respectively to see if the movement axis was “more asymmetrical” or “less asymmetrical” respectively.

### Motility orientation alignment & movement axis analysis for bilateral symmetry

The generated tracks were analyzed alongside Z-stacks of designated Anthrobot from a confocal microscope to see if their morphology was connected to their movement. ImageJ was used to compile the slices of the Anthrobot so that a 3D model could be generated and rotated to render a transformation that visually matches a random frame of the Anthrobot from the timelapse. This random selection could be done as the Anthrobot, despite moving around, did not tend to roll and therefore generally maintained the same orientation throughout a timelapse. Additionally, they often moved with a specific side that always faced forward that was designated as a heading. To see if biases in cilia patterns to one side or lack thereof on an Anthrobot affected its movement this heading would serve as the axis along which a plane of symmetry would be extended to bisect the bot. This plane would be defined by 3 points on the bot along this axis, one placed at the centermost point of the axis within the bot, one on the part of the bot that most visually served as the heading in the video and one on the opposite point of the bot from the heading.

### Neuronal Traversal and Healing Tissue Density Analysis

This procedure was carried out on 30+ files and yielded 20 usable datasets, which were whittled down to 17 after a manual check of tracking quality and excluding videos where bots never touched the wound wall. Using the coordinates of the wound walls and the tracking of the bot, we used the Rbioformats (version 0.0.74)^8^ and Rvision (version 0.6.2)^9^ package tools to test (1) whether bots are more in contact with the wound when they have a higher rotational tendency and (2) when moving on tissue, whether faster bots tend to cover more area i.e., explore better. The lm() function was used to model the data after calculation of proportion of bot on tissue, instantaneous angular velocity and linear speed as variables. Before each model was approved, diagnostics were run on the model using the DHARMa package which included analysis of the residuals.

Afterwards, in order to take a better look at the nuances of interactions between bots and wounds, we decided to limit the data and remove any videos that had bots with very low rotational tendency (<0.33) since they were not stable enough in their rotational behavior. Finally, bots with either very high rotational tendency (>0.7) were removed since they were prone to skidding instead of interacting with the wound walls. Finally, bots whose tracking videos were not optimal i.e they frequently went backward or circularly in the wound were removed since they did not have consistent forward movement that could be correlated with the wound wall. After all these removals, our dataset ended up with 13 examples of wound-bot interactions which could be effectively analyzed. The Gyration was simply the Rotational Tendency values renamed. Our Wound-Trajectory Similarity metric was calculated as the larger absolute value of the correlation between the heading angle of the trajectory and the heading angles of the wound from the surface perpendicular to the bot. These correlation values were then modelled using the lm() function with an expectation of a quadratic relationship for Gyration. The specifics can be seen in the attached code.

Traversal videos of bots moving along a wound within a neuron plate were processed via Adobe Illustrator to see if the bot faithfully followed the edge of the wound. The videos were processed within Adobe Photoshop to generate image sequences compatible with Illustrator for this analysis. The first method of processing aligned the center of the wound at a horizontal line parallel to the bottom of the screen and placed a point on the center of the bot at each frame of the video as well as straight above and below this point on the edges of the wound. Lines were made to connect each respective type of point for both of the edges of the wound and the position of the bot. The output of this for further analysis was a set of coordinates of the end of each line derived from rendering these series of lines as an SVG file and exported as text.

Next, to investigate whether these “bridges” were actually akin to neurons, we decided to analyze the pixel densities of various areas on and surrounding the bridge. In order to prevent confusions regarding this process with regard to intensity of color, we to binarized the image on ImageJ. If the automatic thresholding did not visually appear similar to the raw image, we adjusted the threshold manually. We ended up with six areas of interest: the neurons above the bridge, below the bridge, to the left but adjacent to the left but far, and to the right, both adjacent and far. These areas were defined relative to a FIJI ROI box on the neuronal bridge which tried to encompass the width of the bridge and the height close to the narrowest point of the wound channel that we would interact with. A line of one bridge length or lower if the image size required smaller lines to fit the boxes was used in the vertical and horizontal directions (called hereafter as “bridge length”). The above and below bridge measurements were taken by placing the bounding box one vertical bridge length from the box on the neuronal ridge. The adjacent areas on both sides were defined as 1 horizontal bridge length away from the bridge in the wound. The far areas were 1 bridge length beyond the adjacent areas. The far and adjacent boxes were (vertically) adjusted so they overlaid the wound as much as possible (Figure S6). Finally, we used Analyze>Histogram in FIJI to get the size of the box (which was constant) and the number of pixels of wound tissue (in black) and calculated the proportion. We then used an unpaired two sample T-test with unequal SD to calculate the significance of the difference, if any.

