## Supplementary Information for "Motile Living Biobots Self-Construct from Adult Human Somatic Progenitor Seed Cells"

### Supplemental Figures

#### Supplemental Figure 1

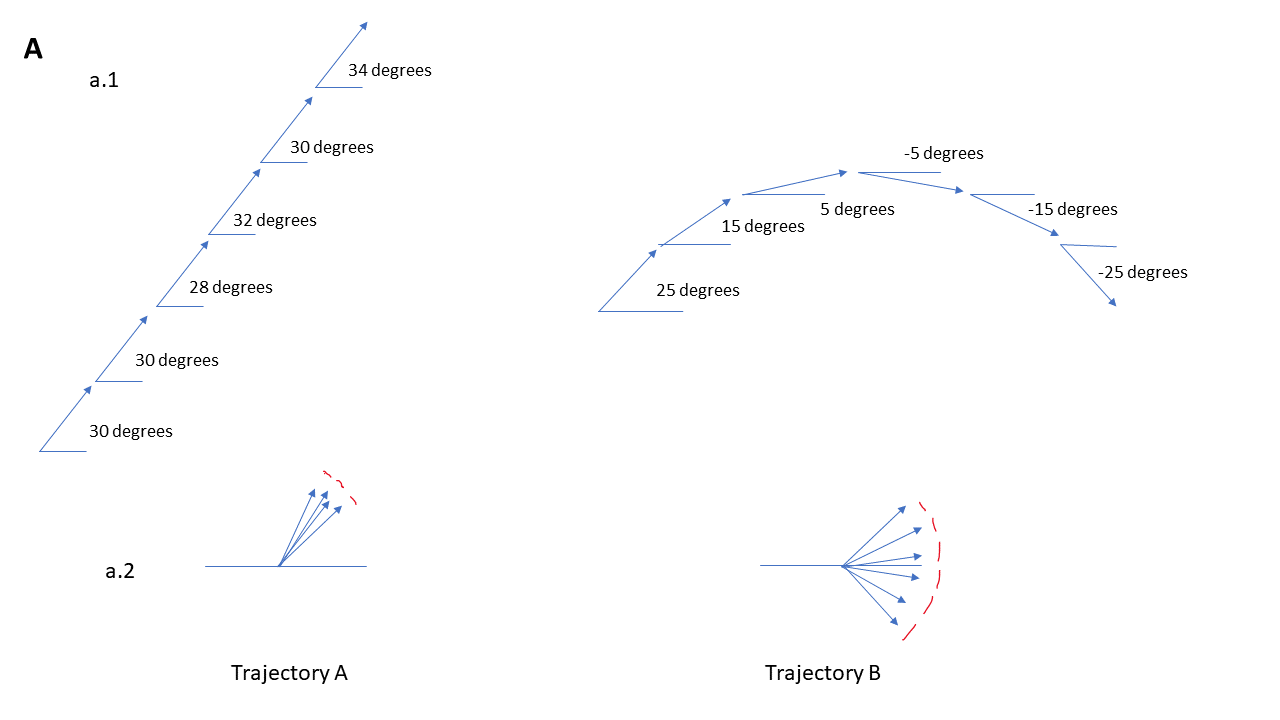

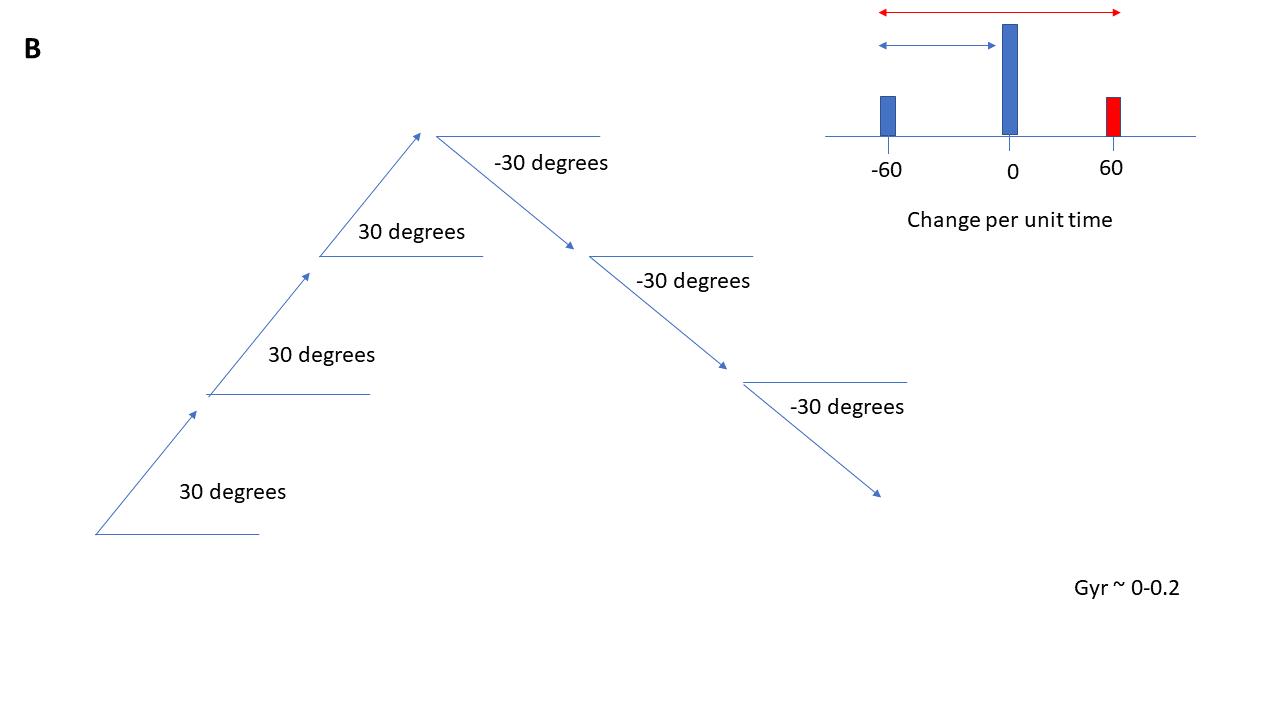

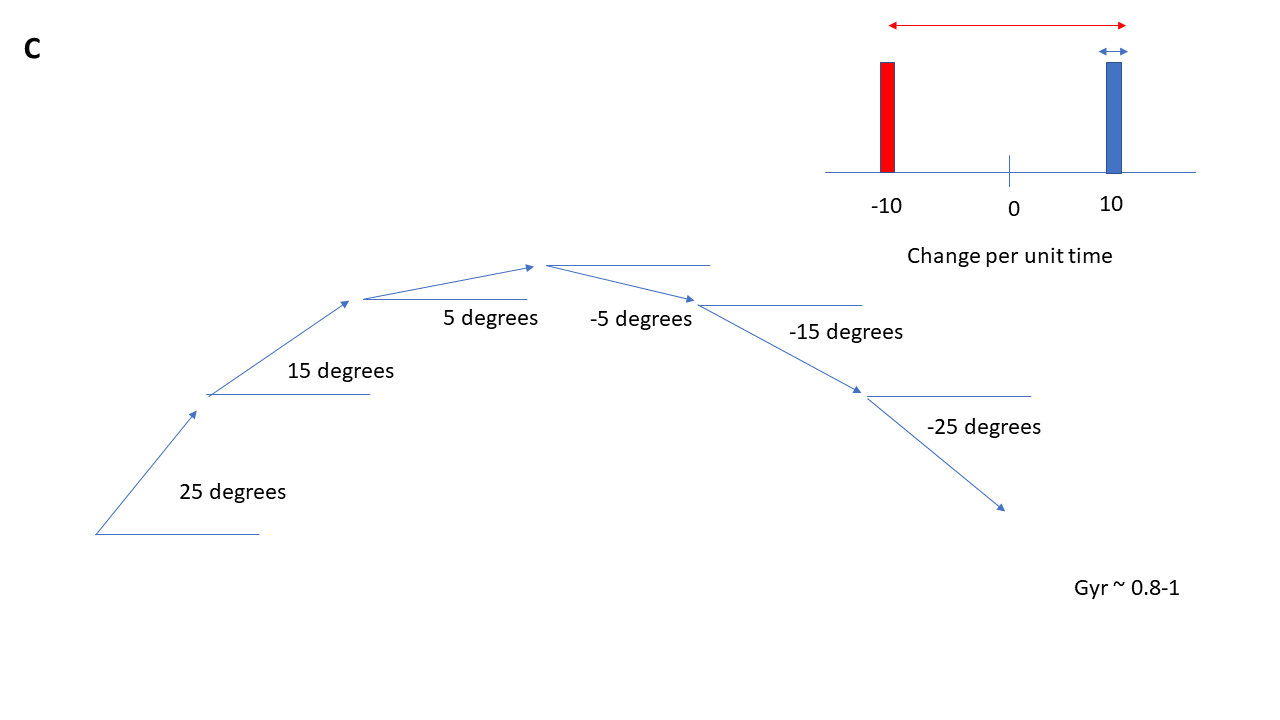

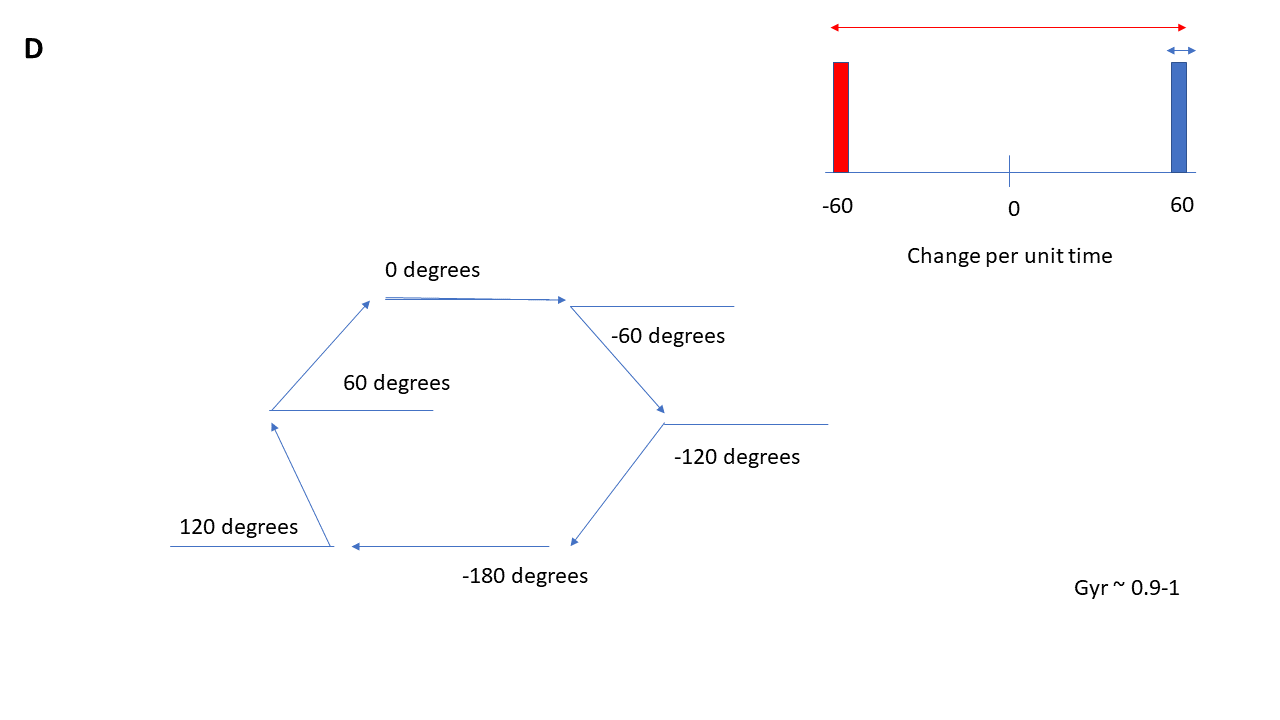

**Figure S1** Sample trajectories that reference a relative x-axis to find heading angle. (A) Visual representation of straightness index, which calculates spread of headings as a whole without taking into account temporal dynamics. (B, C, D) Visual representation of gyration index, which includes the temporal aspect and calculates spread of change in headings relative to their magnitude. Graphically, it is represented as the ratio of the blue length (original circular variance) to the red length (circular variance of the original angular speeds and their additive inverse) on the histograms.

#### Supplemental Figure 2

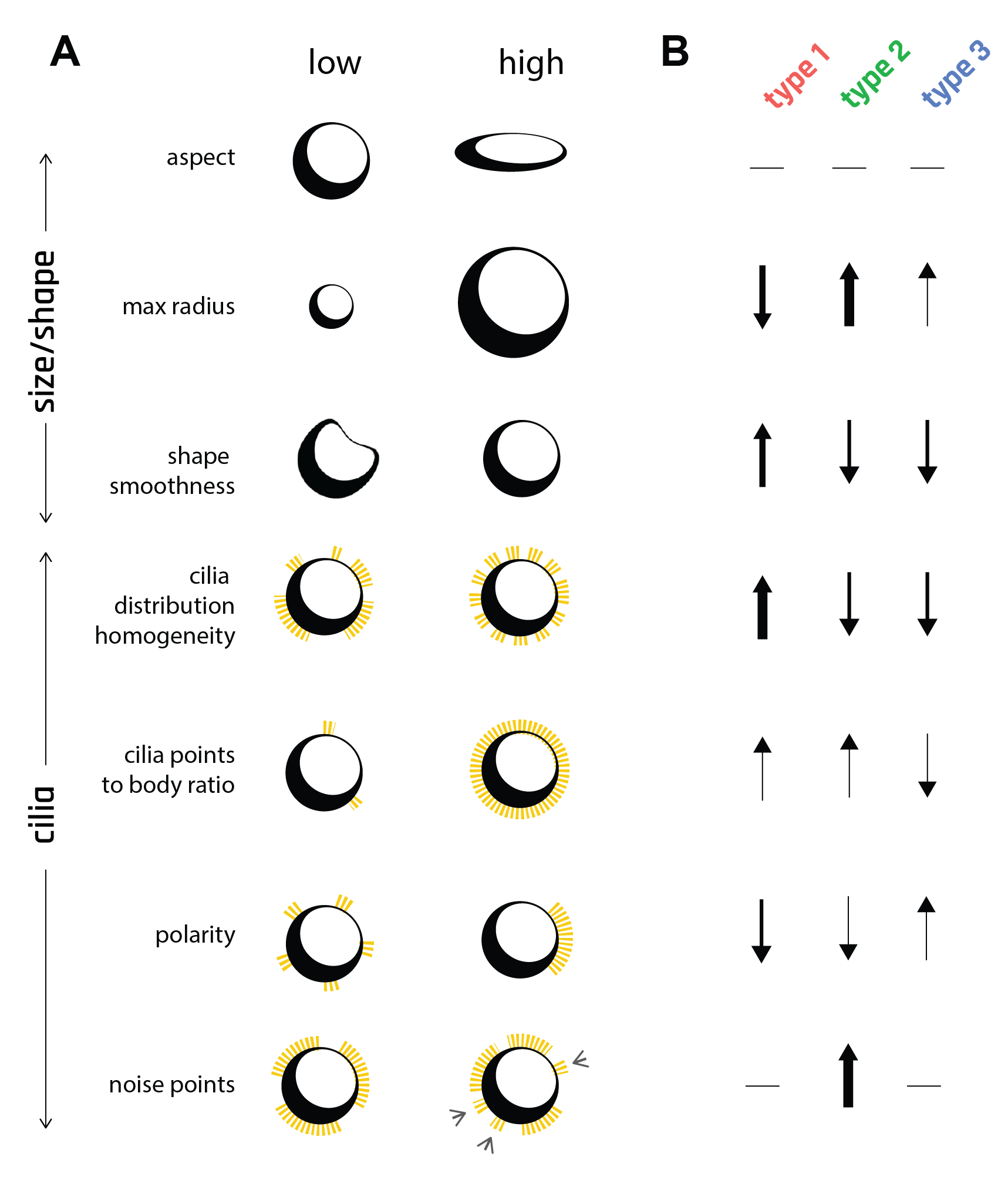

**Figure S2.** Eight morphological indices were used to characterize Anthrobot morphotypes. (A) Visual summary of morphological indices at their extreme points (low, high). (B) Graphic summary of boxplots on Figure 3D, describing 3 different morphotypes. Arrow thicknesses are correlated with the # of standard deviations between a given morphotype’s mean vs the overall population mean for a particular morphological index.

#### Supplemental Figure 3

45 Degrees

|  | Circular | Linear |
| --- | --- | --- |
| Cilia | 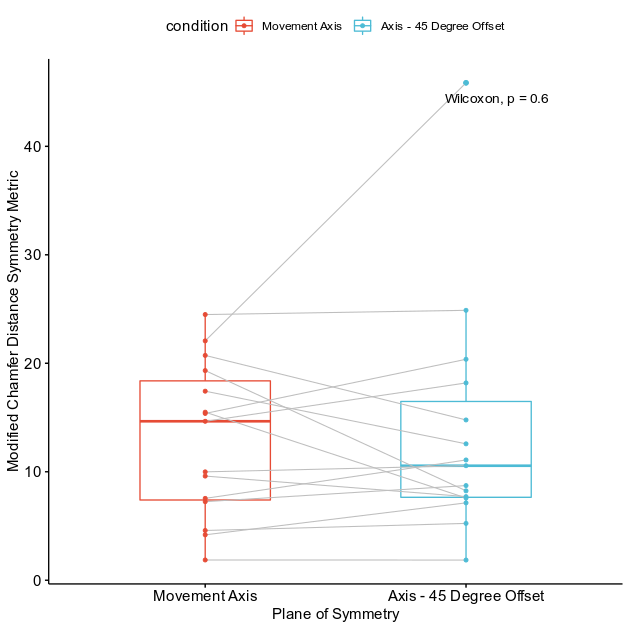 | 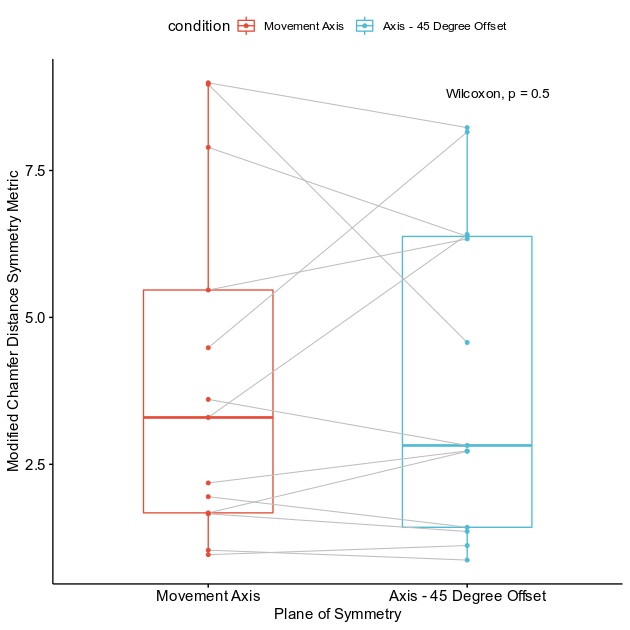 |
| Body | 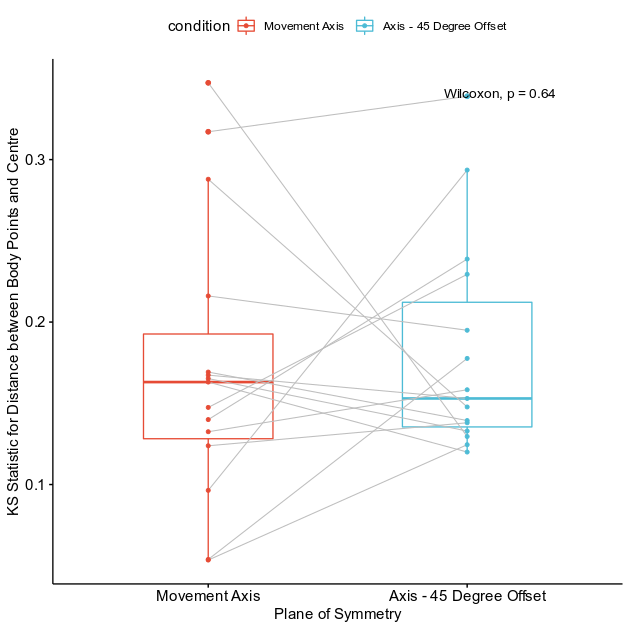 | 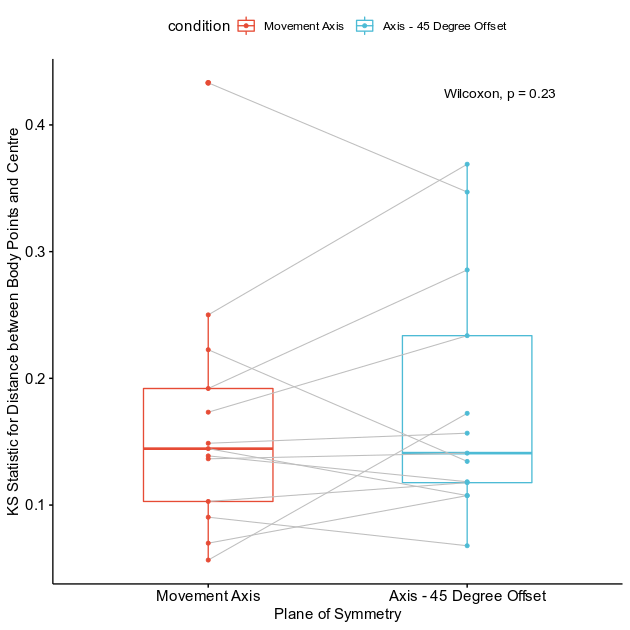 |

90 Degrees

|  | Circular | Linear |
| --- | --- | --- |
| Cilia | 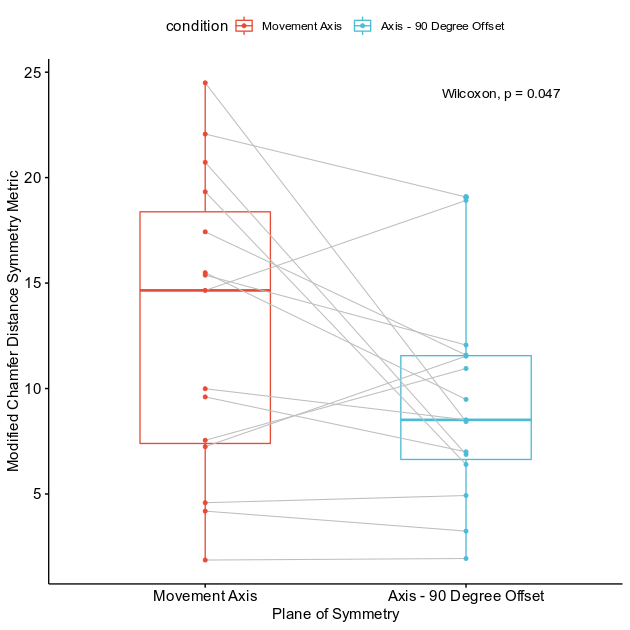 | 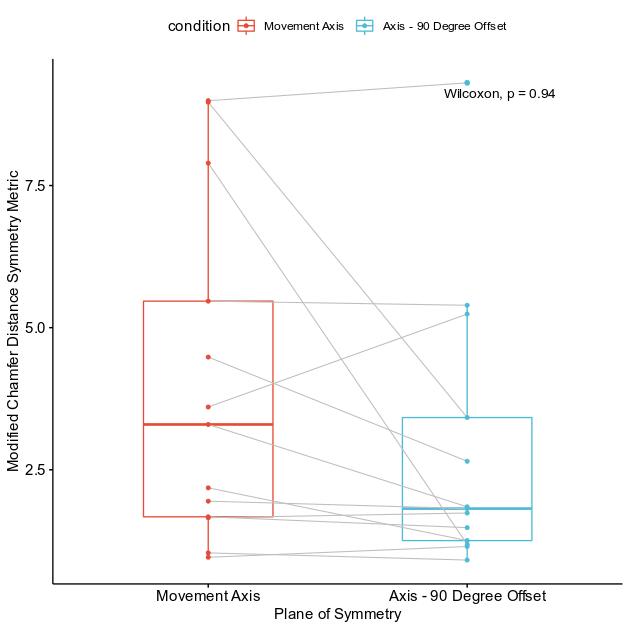 |
| Body | 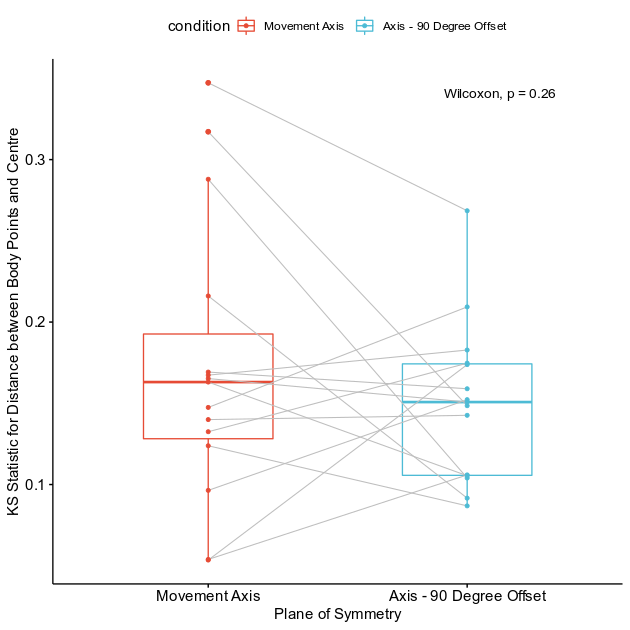 | 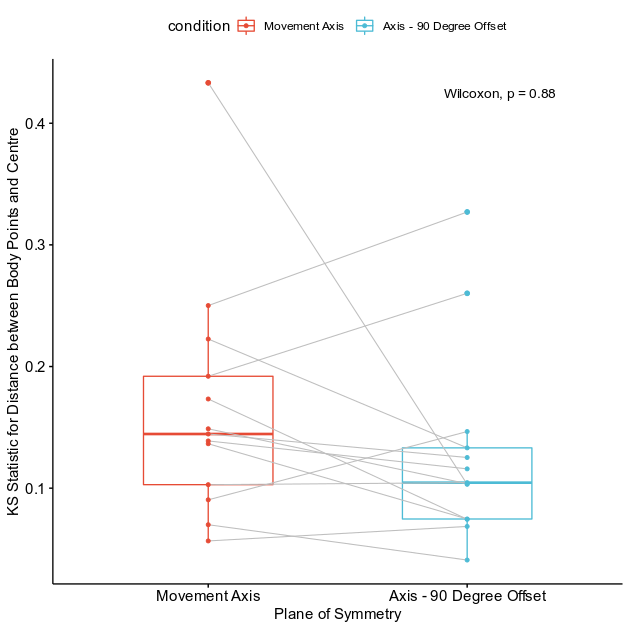 |

135 Degrees

|  | Circular | Linear |
| --- | --- | --- |
| Cilia | 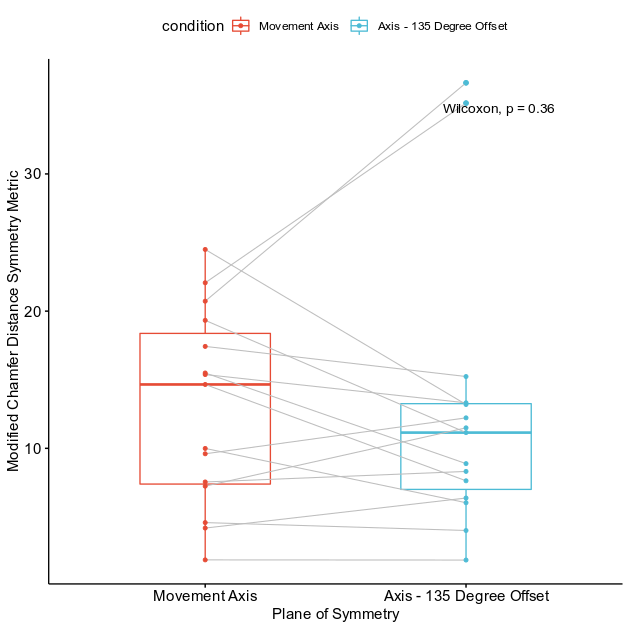 | 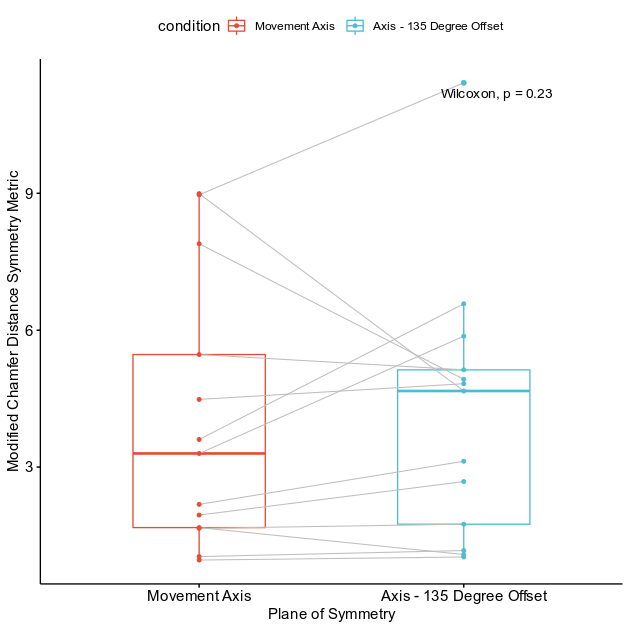 |
| Body | 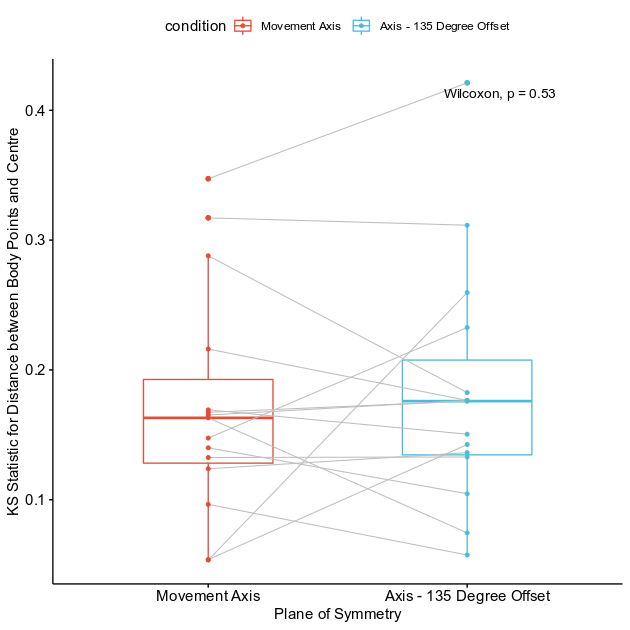 | 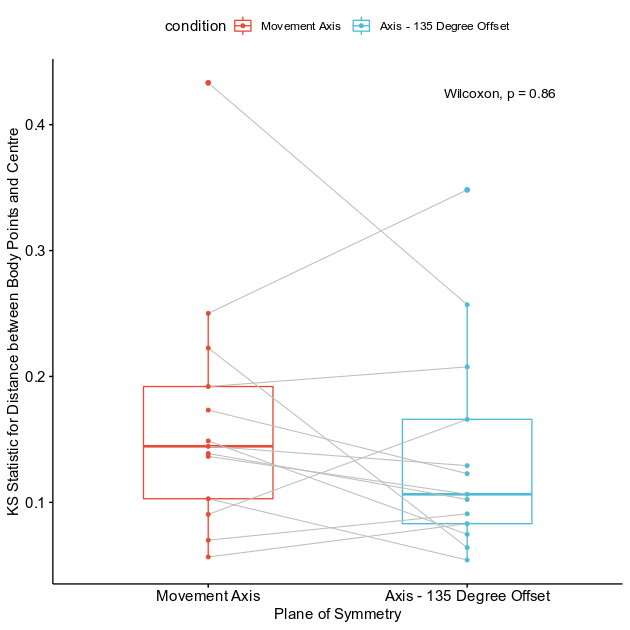 |

**Figure S3** Difference in asymmetry of cilia distribution and body shape between linear and circular bots in respect to the movement axis and its 45, 90, and 135-degree offset axes.

#### Supplemental Figure 4

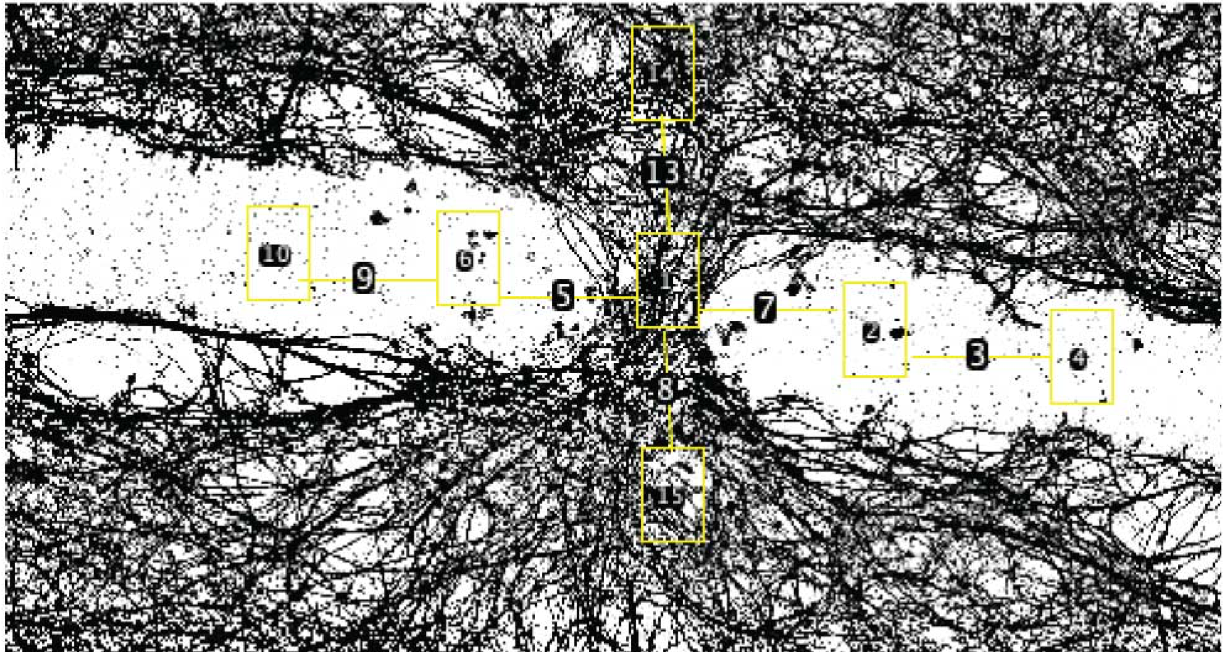

**Figure S4** Sample neuronal density sampling region. Each rectangle represents an area sampled and the lines are consistently the same length, the “bridge length.”

### Supplemental Methods Information
